# Complementary human gene interaction maps from radiation hybrids and CRISPRi

**DOI:** 10.1101/2024.02.05.579036

**Authors:** Desmond J. Smith

## Abstract

The only comprehensive human genetic interaction map was constructed using increased gene copy numbers in radiation hybrid (RH) cells. Recently, a second map restricted to essential genes was created using CRISPRi-induced loss-of-function alleles. Here, the two maps are compared to understand their similarities and differences. Both maps showed significant overlap with protein-protein interaction databases and identified a shared set of interacting genes, although the specific gene pairs differed between approaches. Notably, the RH map exhibited strong overlap with genome-wide association study (GWAS) networks, while the CRISPRi map did not. These findings demonstrate how gain- and loss-of-function alleles reveal distinct yet complementary genetic interaction landscapes.

**NEW & NOTEWORTHY:** This study compared two mammalian genetic interaction networks for cell growth: one using extra gene copies (RH) and another using partial gene suppression (CRISPRi). Both networks overlapped with protein-protein interaction data and identified common interacting genes, yet specific gene pair interactions differed dramatically. Only the RH network predicted GWAS networks. As the first comparison of large-scale mammalian genetic interaction networks, this work reveals how gain- and loss-of-function capture distinct, complementary biological landscapes.

## INTRODUCTION

Constructing comprehensive genetic interaction maps of the human genome has been challenging due to the vast number of potential interactions. With ∼20,000 protein coding genes, there are roughly 2.0 × 10^8^ possible pairwise interactions (∼square of gene number) (1). Inclusion of non-coding transcripts triples the gene count and increases potential interactions to ∼1.8 × 10^9^.

Radiation hybrid (RH) mapping emerged in the 1990s as a highresolution statistical approach for mapping the human genome (2, 3). Human cells are lethally irradiated, fragmenting DNA into short pieces. Cell fusion then transfers random DNA fragments into living recipient cells (Fig. 1*A*). These short fragments enable high-resolution genetic mapping. Markers located close together in the genome are rarely separated by radiation and tend to be co-inherited in RH clones.

**Figure 1.**
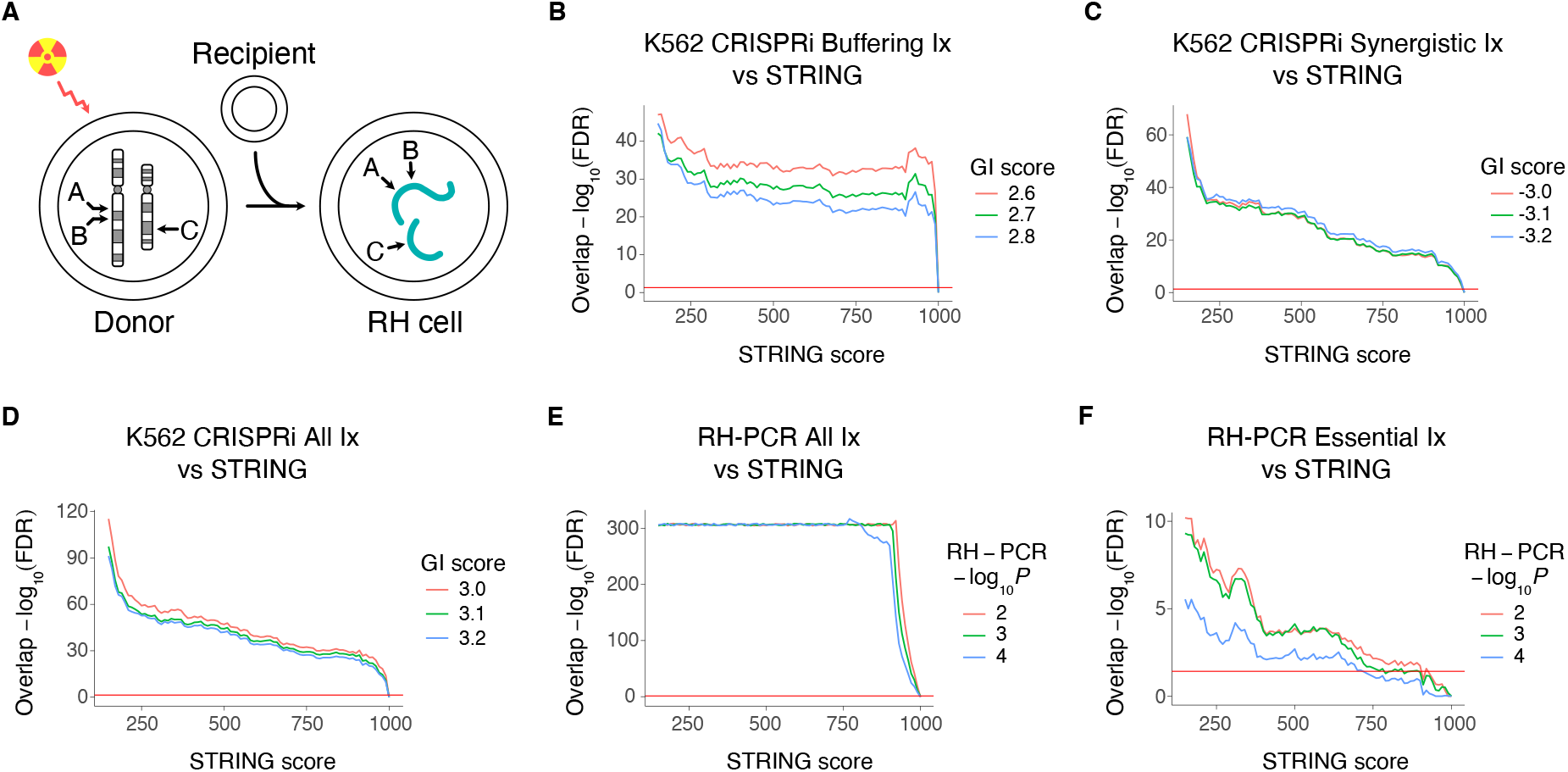
RH-PCR, CRISPRi and STRING interaction networks. *A*: Construction of RH cells. Genes A and B reside close to each other in the genome and tend to be co-inherited in the RH cells. Gene C is a long way from A and B, and is inherited independently unless it interacts with either A or B. Small DNA fragments, torquoise strands. Recipient cell chromosomes not shown. *B*: Overlap of K562 buffering CRISPRi interactions and STRING. Horizontal red line, FDR (false discovery rate) = 0.05. *C*: Overlap of K562 synergistic CRISPRi interactions and STRING. *D*: Overlap of K562 combined buffering and synergistic CRISPRi interactions (All) and STRING. *E*: Overlap of protein coding RH-PCR network (All) and STRING. *F*: Overlap of RH-PCR interactions restricted to essential genes in the CRISPRi dataset and STRING. The higher the STRING score threshold, the greater the confidence in the protein-protein interaction. GI, CRISPRi genetic interaction. Positive GI scores, buffering interactions; negative, synergistic. Ix, interactions.

**Figure 2.**
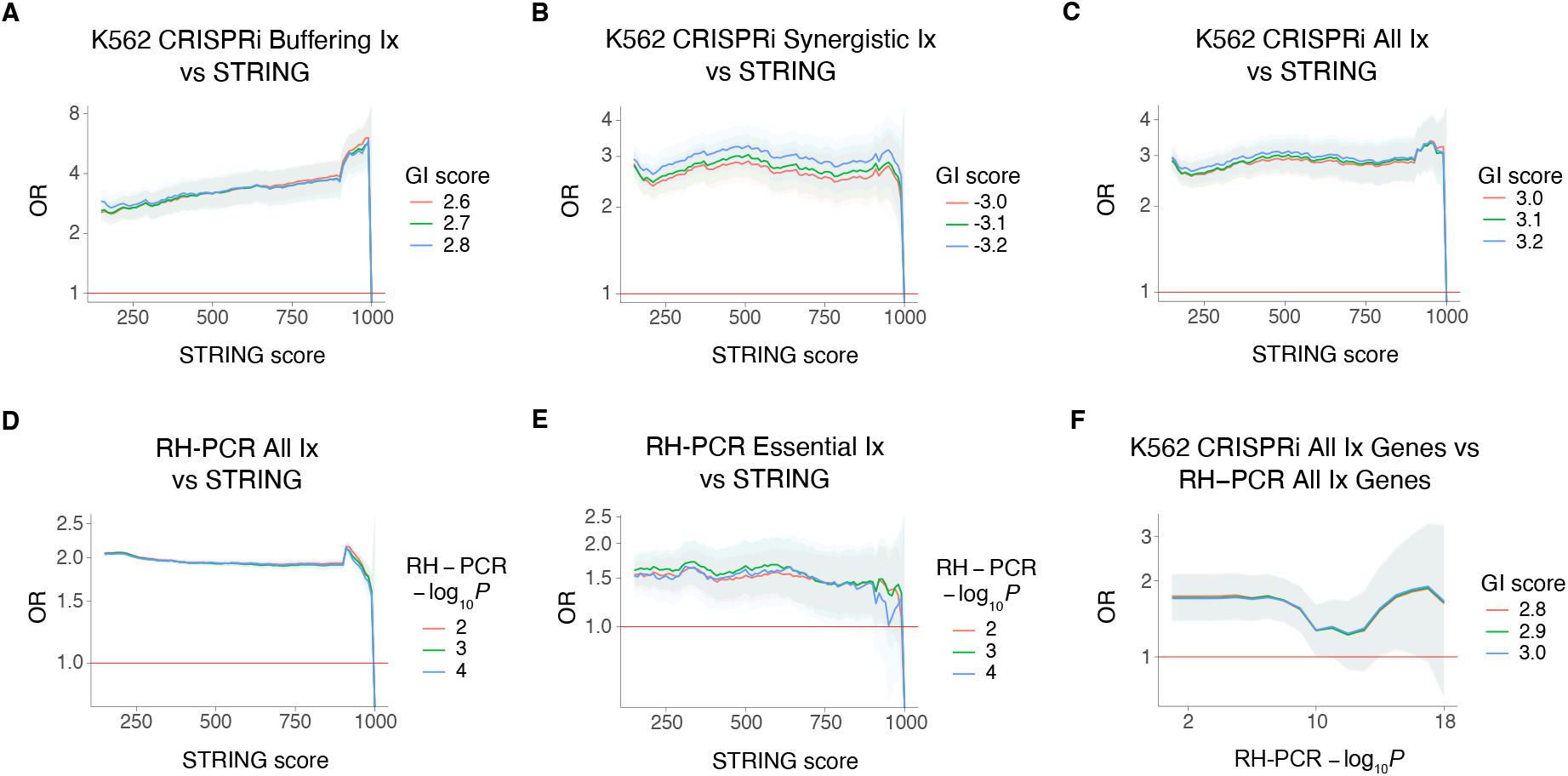
Odds ratios (ORs) for overlaps of RH-PCR, CRISPRi and STRING networks. *A*: K562 buffering CRISPRi interactions and STRING. *B*: K562 synergistic CRISPRi interactions and STRING. *C*: K562 combined buffering and synergistic CRISPRi interactions and STRING. *D*: Protein coding RH-PCR network (All) and STRING. *E*: RH-PCR interactions restricted to essential genes in the CRISPRi dataset and STRING. *F*: Genes involved in K562 combined CRISPRi and protein coding RH-PCR networks. Odds ratios on log_10_ scale. Grey areas, 95% confidence limits. Horizontal red lines, odds ratio = 1.

Distant genes typically segregate independently. However, interacting genes show co-inheritance even when genomically distant (4). For example, when an extra copy of one gene’s toxic effect on cell growth is blocked by an extra copy of a distant gene, these genes show significant co-inheritance in RH clones – an “attractive” interaction. Conversely, two distant genes that together inhibit proliferation show significant lack of co-inheritance – a “repulsive” interaction.

Each RH clone harbors ∼10% to 25% of the genome as extra copies, corresponding to ∼2,000–5,000 protein coding genes. This high multiplexing provides extraordinary efficiency, enabling evaluation of all 1.8 × 10^9^ potential interactions across coding and non-coding genes.

We constructed a single gene resolution interaction network for the human genome using the RH approach (4). PCR genotyping of 560 RH clones from six panels (mouse, rat, dog, and three human) yielded interaction networks that each significantly overlapped when aligned to the human genome (*P* = 1.9 × 10^−11^), allowing dataset combination for increased statistical power. Recipient cells were A23 cells (male Chinese hamster lung fibroblasts) for 472 clones (84%), while HTK3-cells (female Chinese hamster ovary epithelial cells) were used for the 88 dog clones (16%).

Single gene resolution of the RH-PCR network was confirmed by its optimal overlap with protein-protein interaction networks at nearest gene resolution (odds ratio, OR > 10^21^) and narrow interaction peaks (< 20 kb −2log_10_ *P* intervals) (4). The RH network specifically excluded gene pairs within the maximum fragment length (10 Mb), eliminating genomic linkage as a confound.

Among significant protein coding RH-PCR interactions, 99.97% were attractive. This preponderance is unlikely to reflect methodological bias, as ∼96% of marker pairs had equivalent statistical power to detect attraction or repulsion (4). The imbalance suggests that extra copies of distant genes promoting growth (attractive interactions) occur more frequently than extra copies synergistically inhibiting cells (repulsive interactions).

Other technologies can assess genetic interactions in mammalian cells, including RNA interference (RNAi) for gene knockdowns (5), CRISPR for knockout (null) alleles (6), CRISPR interference (CRISPRi) for partial loss-of-function (hypomorphic) alleles (7), and CRISPR activation (CRISPRa) for constitutive overexpression (8).

These approaches can probe single gene mutant interactions with nearly complete arrays of protein coding genes. However, evaluating all pairwise combinations remains infeasible due to the vast numbers involved. Unlike RH approaches that multiplex thousands of genes per cell, CRISPR and RNAi methods evaluate one gene pair in each cell.

Nevertheless, CRISPR and RNAi technologies have explored small interaction subsets, including ∼2.6 × 10^3^ cancer gene combinations across three cell lines (9), ∼1.0 × 10^5^ combinations using orthogonal CRISPR/CRISPRa systems (10), and ∼5.7 × 10^3^ to 1.1 × 10^4^ combinations of chromatin regulating gene via CRISPRi or RNAi, revealing that significant genetic interactions predicted corresponding protein associations (11, 12). While simultaneous multigene CRISPR targeting is being developed (13, 14), current efficiencies limit targeting to ∼6-7 genes – far below the thousands in RH cells.

Recently, CRISPRi was used to examine interactions among ∼1.0 × 10^5^ pairwise combinations of ∼440 protein coding genes (7). CRISPRi uses catalytically inactive Cas9 (dCas9) fused to KRAB repressor to silence genes and create partial loss-of-function alleles. Targeted genes show maximal silencing, but adjacent genes may also be repressed. The screen used K562 cells (human chronic myelogenous leukemia) and Jurkat cells (human T-cell leukemia) stably expressing dCas9-KRAB, transduced with a library of lentiviruses, each delivering two sgRNAs to target a gene pair in a cell.

The CRISPRi study evaluated genes with moderate individual loss-of-function growth effects, as essential gene silencing causes cell death. Genes evaluated in the CRISPRi study are termed here “essential” for brevity. The proteins encoded by the genes in the CRISPRi screen resided in the major cellular compartments and participated in diverse cellular activities. The CRISPRi network assessed ∼0.05% of all possible protein coding interactions, contrasting with the genome-wide RH-PCR network.

The CRISPRi screen identified two interaction types: buffering (combined loss-of-function alleles yield better growth than their sum – statistically a positive interaction) and synergistic (combined alleles yield poorer growth – statistically negative). Note that “buffering” can also describe weaker-than-expected deletion phenotypes due to paralog compensation (15, 16).

Here, the RH-PCR and CRISPRi networks – the largest independent mammalian genetic interaction datasets to date – are compared. The investigation reveals the contrasts between networks constructed using extra copy and partial loss-of-function variants and their connection with other biological networks, including protein-protein (17) and genome-wide association study (GWAS) networks (18).

## MATERIALS AND METHODS

### Data Analysis

Statistical analyses were performed using R (19). Logistic regression employed `biglm` (20). Graphs were generated using `ggplot2` and `cowplot` (21, 22). Receiver operating characteristic (ROC) curves and areas under the curve (AUCs) were calculated using `pROC, plotROC`, and `caTools` (23, 24, 25). Precision-recall (PRC) analyses employed `PRROC` (26). Networks were visualized using Cytoscape in the R environment via `RCy3` (27, 28).

### CRISPRi Network

The CRISPRi knockdown screen used two replicates of K562 and Jurkat cells that constitutively expressed dCas9-KRAB. Cells were transduced with a lentiviral library in which each virus delivered two sgRNAs targeting a gene pair. Cell growth was quantified at day 5 post-infection (∼10 population doublings) by sequencing three lentiviral barcodes (19, 19, and 38 bp) on an Illumina HiSeq 2500 or 4000 with ∼509–931 million (M) reads (7).

Buffering or synergistic CRISPRi interactions were classified as significant if the average genetic interaction (GI) score of the two replicates was greater or less than the 95% confidence intervals of the negative controls, respectively. For K562 cells, the GI threshold was 2.5 for buffering interactions and −2.6 for synergistic. For Jurkat cells, the thresholds were 1.9 and −2.0, respectively. Combined CRISPRi interactions were thresholded on highest absolute GI score, 2.6 for K562 and 2.0 for Jurkat.

Genes targeted in the CRISPRi network were protein coding with moderate loss-of-function growth effects, listed in (7) and the repository noted in **DATA AVAILABILITY**.

### STRING Network

The STRING dataset (v11) contains 5,879,727 human proteinprotein interactions compiled from multiple publicly available sources (17). Each interaction is assigned a confidence score ranging from 150 (lowest) to 999 (highest).

### GWAS Network

A publicly available dataset summarizing published GWAS results was employed (18). Reflecting the high mapping resolution of modern GWASs, candidate genes were typically identified as single genes nearest to the most significant variants. Of 516,161 entries in the GWAS dataset, 317,224 (61%) were single gene entries. The remaining 198,937 entries (39%) consisted of multigene intervals bounded by two candidate genes.

Both single and multigene entries were retained in the GWAS dataset to capture uncertainties in gene assignment and avoid overfitting. For multigene entries, the gene at the interval start was selected, except for 10,168 intervals (5%) lacking a nominated start gene, for which the end gene was used. The cleaned GWAS dataset comprised 30,211 unique genes underlying 25,461 traits.

The GWAS network was constructed by linking all genes associated with the same phenotype. Interaction −log_10_ *P* values were calculated as the mean of each trait. To enable comparison with analogous RH-PCR networks, two additional GWAS networks were constructed: one with at least one gene of each interaction on the X but not Y chromosome, and one consisting of essential genes plus genes one link away.

Overlap comparisons of essential RH-PCR and CRISPRi networks with GWAS used the non-downsampled GWAS networks. However, comparisons involving X chromosome-specific RH-PCR, essential RH-PCR plus one, and full protein coding RH-PCR networks used corresponding GWAS networks downsampled 20-fold to enable manageable computation times.

To evaluate whether linkage substantially confounded the GWAS network, we created a trimmed dataset containing only single gene entries. The protein coding RH-PCR network was compared with the single gene GWAS network downsampled 20-fold. Comparisons with CRISPRi networks used the complete single gene GWAS network.

Overlaps between the protein coding RH-PCR network and single trait GWAS subnetworks used non-downsampled subnetworks, except for height which was downsampled twofold.

### Overlap Analyses

Network overlaps were evaluated as described (29), except that Fisher’s Exact Test was used instead of the *χ*^2^ approximation when expected values in all table cells exceeded 50. Numbers used for significance testing of optimal or complete network overlaps are provided in Supplemental Tables S1–S9. Total population size depended on the specific comparison.

Overlaps of CRISPRi with STRING protein-protein interactions, essential RH-PCR, and GWAS networks used the number of interactions tested in the CRISPRi study, depending on cell line (K562 or Jurkat) (Supplemental Tables S1, S3, S6). The protein coding RH-PCR network and RH-PCR networks randomly downsampled to match CRISPRi network size were compared with STRING, 20-fold downsampled GWAS, and single trait GWAS networks using all pairwise non-redundant combinations of protein coding genes in `GENCODE` v31 (30). The total interaction number was 199,490,325 (Supplemental Tables S2, S5, S7, S8).

Overlaps between STRING and the protein coding RH-PCR network randomly downsampled to match CRISPRi network size used 20 independent downsamples to calculate empirical standard deviations (s.d.) of the odds ratio (OR) (Supplemental Table S2).

Comparison of the RH-PCR essential plus one and 20-fold downsampled GWAS networks used all non-redundant pairwise combinations of unique genes in the RH-PCR essential plus one network and the analogous downsampled GWAS network (84,337,578 interactions) (Supplemental Table S5).

To assess reproducibility of overlaps using the downsampled GWAS network, the empirical s.d. of the OR at optimal overlap was evaluated using 20 independent downsamples (Supplemental Table S5). Standard deviations derived from downsamples were consistent with significant overlaps. The twofold downsampled height GWAS subnetwork was independently replicated to provide an s.d. of the OR (Supplemental Table S7).

RH-PCR networks with one gene restricted to X or Y chromosomes were compared with STRING using a population equal to the non-redundant pairwise combinations of protein coding genes on the relevant sex chromosome and autosomes, plus pairwise combinations of genes on the relevant sex chromosome (RH-PCR chrX: 16,560,975 interactions; RH-PCR chrY: 1,260,039 interactions) (Supplemental Table S2). Gene overlap between CRISPRi and RH-PCR networks used a population equal to the number of unique protein coding genes in `GENCODE` (19,942 genes) (Supplemental Table S4).

Potential confounds of gene expression and evolutionary conservation were controlled by including them as covariates in overlap analyses using logistic regression (Supplemental Table S9). Gene expression in K562 and Jurkat cell lines used RNA-Seq datasets (31, 32), while expression of genes in the RH-PCR network used microarray data from a human RH panel (33). Evolutionary conservation was assessed using phyletic conservation – the number of species with orthologs of the interacting genes (1, 16, 31, 32, 34). Since centrality reflects the likelihood of forming an interaction, these measures were omitted as covariates due to collider bias.

### Growth Promoting and Suppressing Genes

To examine relationships between gene essentiality and network centrality in the protein coding RH-PCR and STRING networks, data was used from a single gene CRISPR knockout study in K562 cells (34). Null alleles were created by transducing a lentiviral library in which each virus targeted a single gene using an sgRNA. Effects on cell proliferation (∼14 doublings) were evaluated by quantifying sgRNA barcode abundance using 50–75 M reads on an Illumina HiSeq 2500 sequencer. Growth genes significantly decreased proliferation when deleted, while growth suppressing genes increased proliferation (adjusted *P* < 0.05).

### Gene Expression

The connection between gene expression and genetic interactions was evaluated. RNA-Seq of K562 cells used ∼130 M paired-end reads of 76 bp on an Illumina Genome Analyzer IIx (31), while RNA-Seq of Jurkat cells used 44 M paired-end reads of 90 bp on an Illumina HiSeq 2000 machine (32). Expression data for the RH-PCR network employed a microarray dataset from a human RH panel obtained using Illumina HumanRef-8 v1.0 BeadChips (33).

### Clustering of RH-PCR Network

Clustering of interactions restricted to CRISPRi essential genes in the RH-PCR network used Euclidean distance measures in heatmap.2 (35). Clusters were defined using dendrogram cuts closest to the origin, and genes assigned to gene ontology (GO) categories using ontologyIndex (36, 37). Significance values for GO enrichment of clusters were calculated using Fisher’s Exact Test. Cell compartment assignments in the essential RH-PCR network were performed as described (7).

## RESULTS

### Properties of RH-PCR and CRISPRi Networks

The complete RH-PCR network contained 7,248,479 significant interactions involving 18,220 genes (false discovery rate, FDR < 0.05) (4, 38). When restricted to protein coding genes, the network comprised 2,705,945 significant interactions among 10,897 genes, representing 1.4% of all possible interactions. Among these protein coding interactions, 912 (0.03%) were repulsive and the remainder attractive. None of the genes participating in repulsive interactions appeared in the CRISPRi dataset.

CRISPRi interaction strengths were quantified using genetic interaction (GI) scores (7). Buffering interactions received positive GI scores, indicating that cells with two partial loss-of-function alleles grew better than expected from their additive inhibitory effects. Higher positive scores indicate stronger buffering interactions.

Conversely, synergistic interactions received negative GI scores, indicating that cells grew more poorly than expected from additive effects. More negative scores indicate stronger synergistic interactions. Beyond examining buffering and synergistic interactions separately, their combined properties were analyzed by considering all significant interactions regardless of sign.

Significance thresholds for GI scores, established using negative controls, were > 2.5 for buffering and < −2.6 for synergistic interactions in K562 cells and > 1.9 and < −2.0 in Jurkat cells. For combined networks, absolute value thresholds were 2.6 for K562 and 2.0 for Jurkat.

In K562 cells, CRISPRi testing of 100,128 potential interactions among 448 essential genes identified 3,885 (3.9%) significant interactions involving 446 genes. Of these, 1,493were buffering and 2,392 synergistic. In Jurkat cells, testing 95,266 interactions among 436 essential genes revealed 2,552 (2.7%) significant interactions involving 424 genes, with 1,318 buffering and 1,234 synergistic. Overall, 426 genes and 104,443 interactions were tested in both cell lines. The protein coding RH-PCR network was substantially larger than the CRISPRi network, containing ∼25 times more genes and ∼840 times more significant interactions.

### Network Subsets

Six subsets of the complete RH-PCR network were used for network comparisons:

1. The protein coding RH-PCR network (“All”).
2. The protein coding RH-PCR network restricted to essential genes evaluated in the CRISPRi study. This “essential” RH-PCR network contains 279 genes and 1,778 interactions for K562, or 273 genes and 1,686 interactions for Jurkat. When combined for genes tested in either cell line, the network comprises 285 genes and 1,839 interactions, enabling equally powered overlap comparisons with the CRISPRi networks.
3. The essential RH-PCR network expanded to include genes linked by one interaction (“essential plus one”). This network contains 10,497 genes and 140,308 interactions for K562, or 10,497 genes and 140,216 interactions for Jurkat. The combined network comprises 10,497 genes and 140,369 interactions, intermediate in size between the first two subsets.
4. The protein coding RH-PCR network randomly downsampled to match the number of interactions evaluated in the CRISPRi study for either K562 or Jurkat cells. Overlaps for this “RH-PCR ds” (downsampled) network used the mean results from 20 independent downsamples. The network contains 8,265 ± 20 s.d. genes and 104,443 interactions, similar in size to the essential plus one network.
5. The protein coding RH-PCR network filtered to include only interactions with at least one gene on the X chromosome but excluding Y chromosome genes. This “RH-PCR chrX” network contains 10,071 genes and 151,980 interactions.
6. The protein coding RH-PCR network filtered to include only interactions with at least one gene on the Y chromosome (excluding X chromosome genes). This “RH-PCR chrY” network contains 3,614 genes and 8,440 interactions.

Four subsets of the CRISPRi networks were analyzed:

1. Buffering interactions, with significant positive GI scores.
2. Synergistic interactions, with significant negative GI scores.
3. Combined interactions, with significant GI scores regardless of sign.
4. Chromosome X-filtered CRISPRi network containing only interactions with at least one gene on chromosome X. In K562 cells, 6,167 potential interactions among 448 essential genes were tested, yielding 339 (5.5%) significant interactions involving 225 genes (88 buffering, 251 synergistic). In Jurkat cells, 5,590 potential interactions among 437 genes were tested, yielding 252 (4.5%) significant interactions involving 167 genes (109 buffering, 143 synergistic). The CRISPRi dataset contained no Y chromosome genes.

For network comparisons involving CRISPRi data, K562 cell statistics are reported unless otherwise noted. Jurkat cell results were essentially similar (**SUPPLEMENTAL MATERIAL**).

### Overlap Significance

The significance of overlap between two genetic networks depends on the interaction thresholds applied to each. Rather than relying on a single threshold, the entire overlap landscape was surveyed by systematically scanning threshold parameters from least to most stringent (29). Overlap significance was calculated at each threshold and corrected for multiple testing using false discovery rates (38). This approach provides a comprehensive view of the overlap landscape rather than a single snapshot at one threshold.

### Overlaps of RH-PCR and CRISPRi Networks with STRING

First to be examined was the overlap of the protein coding RH-PCR and CRISPRi interaction networks with the STRING repository of protein-protein interactions (17).

All three CRISPRi interaction types – buffering, synergistic, and combined – showed significant overlap with STRING in both K562 and Jurkat cells (K562 combined interactions: optimum FDR = 6.2 × 10^−116^, OR = 3.0, Fisher’s Exact Test) (Fig. 1*B-D*, 2*A-C*, Supplemental Table S1, Supplemental Figs. S1-S3). The protein coding RH-PCR network also showed significant overlap with STRING (optimum FDR = 4.3 × 10^−319^, OR = 1.9) (Figs. 1*E*, 2*D*, Supplemental Table S2, Supplemental Fig. S4), but with greater significance than CRISPRi (Fig. 3*A*, Supplemental Fig. S4), reflecting the more comprehensive nature of the RH-PCR network. As expected, overlap significance decreased with increasing threshold stringency due to fewer interactions meeting the criteria (Supplemental Fig. S5).

**Figure 3.**
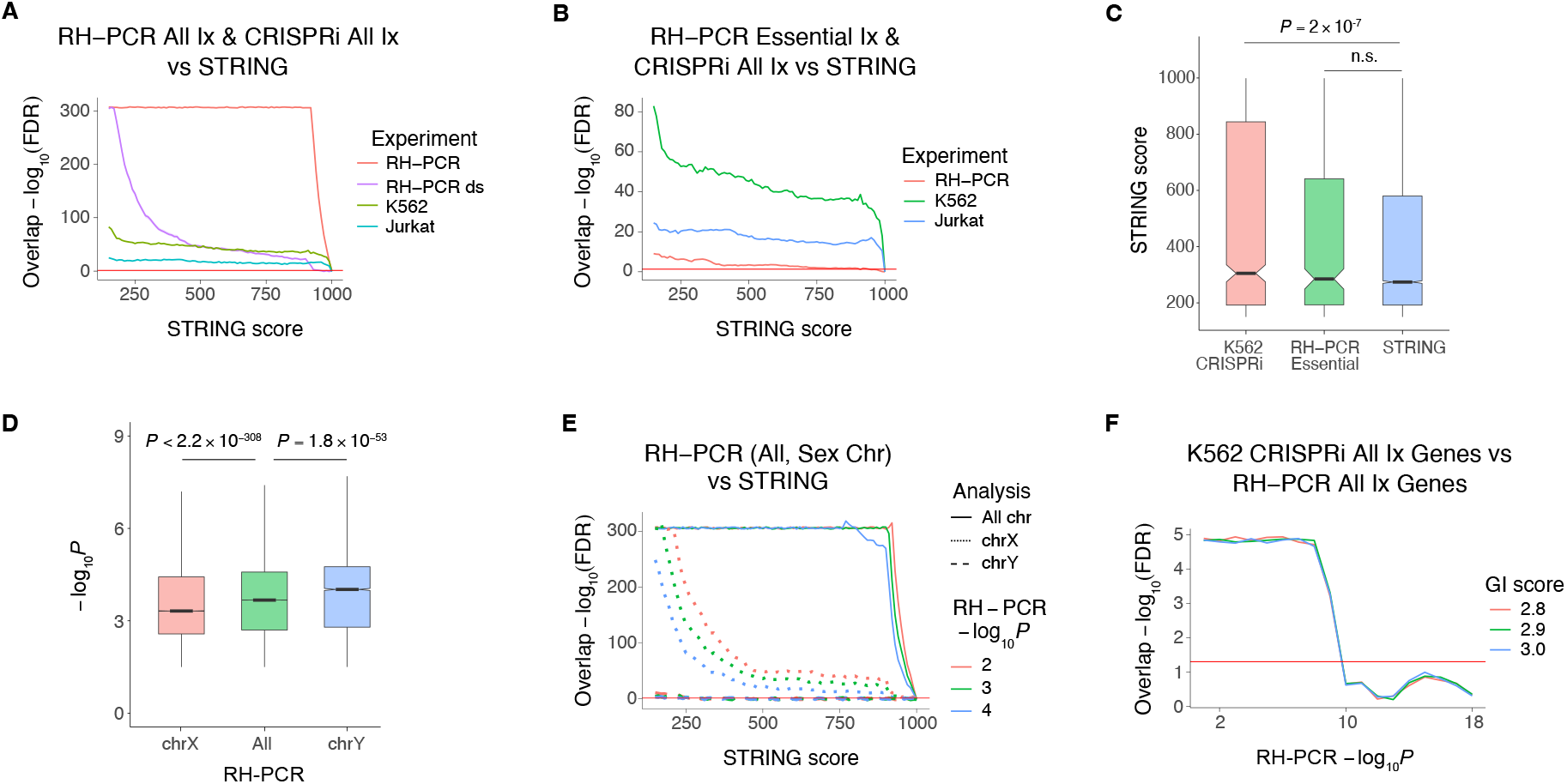
Overlaps of genetic interaction networks and their participating genes. *A*: Overlaps of protein coding RH-PCR network, downsampled RH-PCR network (RH-PCR ds), and K562 and Jurkat combined buffering and synergistic CRISPRi networks with STRING. RH-PCR downsampled network 95% confidence interval shown, but hard to see because narrow. Horizontal red line, FDR (false discovery rate) = 0.05. *B*: Overlaps of essential RH-PCR as well as K562 and Jurkat combined CRISPRi networks with STRING. *C*: STRING scores for interactions specific to K562 combined CRISPRi, essential RH-PCR and STRING networks. *D*: Significance values for the chrX and chrY specific RH-PCR networks and the protein coding RH-PCR network. *E*: Overlaps of the protein coding RH-PCR network and chrX and chrY specific RH-PCR networks with STRING. *F*: Overlaps of genes involved in interactions in K562 combined CRISPRi and protein coding RH-PCR networks.

When restricted to the essential genes evaluated in the CRISPRi study, RH-PCR interactions still showed significant overlap with STRING (optimum FDR = 4.7 × 10^−10^, OR = 1.5) (Figs. 1*F*, 2*E*, Supplemental Table S2, Supplemental Fig. S4), though with less significance than CRISPRi (Fig. 3*B*, Supplemental Fig. S4).

Randomly downsampling the RH-PCR network to match the number of interactions tested in the CRISPRi study did not reduce the optimum overlap with STRING compared to the full protein coding RH-PCR network (optimum FDR = 7.4 × 10^−321^, OR = 2.1 ± 0.03 s.d.), and remained more significant than CRISPRi network overlaps with STRING (Fig. 3*A*, Supplemental Table S2). However, the downsampled network’s smaller size caused overlap significance to decay more rapidly than the full protein coding network as stringency increased.

To control for potential confounds, logistic regression with gene expression and evolutionary conservation as covariates was used to evaluate overlaps among the CRISPRi, RH-PCR, and STRING networks. This analysis yielded similar overlap results to Fisher’s Exact Test (Supplemental Fig. S6).

Interactions in the K562 or Jurkat CRISPRi networks exhibited significantly higher STRING scores than null expectations (K562: *t*[1,1256] = 5.2, *P* = 2.3 × 10^−7^; Jurkat: *t*[1,685] = 4.9, *P* = 1.5 × 10^−6^; Welch Two Sample t-test) (Fig. 3*C*). In contrast, interactions in the essential RH-PCR network showed no significant difference from null. This suggests that RH-PCR interactions more broadly represent protein-protein interactions, whereas CRISPRi interactions preferentially capture those with higher STRING scores.

### Sex Chromosome-Specific RH-PCR Networks

Male recipient cells were used in 84% of RH-PCR clones. In these cells, retained X or Y chromosome genes exhibit a 2-fold copy number change (two copies versus one), compared to autosomal genes which show a 1.5-fold change (three copies versus two). Even in the 16% of clones using female recipient cells, retained X chromosome genes show a functional 2-fold increase due to random inactivation of recipient X chromosomes.

The larger relative copy number increase for sex chromosome genes might be expected to produce stronger genetic interactions than autosomes. However, transcript profiling of RH panels has revealed an additional dosage compensation mechanism beyond X chromosome inactivation (33, 39). Retained X chromosome genes in RH cells actually show blunted expression increases compared to autosomes.

This uncoventional dosage compensation cannot be attributed to X chromosome inactivation, as radiation-induced fragmentation separates X chromosome DNA fragments from the inactivation center. However, the dosage effect aligns with observations that human X-linked genes exhibit lower expression than autosomal genes (40). The attenuated expression response to extra X chromosome copies in RH cells may produce weaker genetic interactions.

To determine whether copy number effects or blunted expression increments predominate, the interaction strengths of X and Y chromosome genes were compared to the complete protein coding RH-PCR network (Fig. 3*D*).

X chromosome genes showed significantly weaker interactions than the complete network (*t*[1,169496] = 47.6, *P* < 2.2 × 10^−308^, Welch Two Sample t-test), indicating that unconventional dosage compensation outweighs copy number effects. Conversely, Y chromosome genes showed stronger interactions than the complete network (*t*[1,8478] = 15.5, *P* = 1.8 × 10^−53^, Welch Two Sample t-test), demonstrating significant copy number effects for this chromosome, which lacks dosage compensation. The weaker X chromosome interactions are consistent with the need for greater functional flexibility to accommodate copy number variation between the sexes (40).

The CRISPRi networks showed no significant differences in X chromosome gene interaction strength compared to complete networks (*P* > 0.1), though statistical power was limited by sample size. The CRISPRi networks contained no Y chromosome genes.

Overlaps between X chromosome-specific RH-PCR networks and STRING matched those of the protein coding RH-PCR network (optimum FDR = 3.7 × 10^−316^, OR = 2.1), though overlap significance decayed more rapidly for the X chromosome network due to its smaller size (Fig. 3*E*, Supplemental Table S2). Thus, network overlaps for X chromosome genes were not substantially affected by either copy number or dosage effects.

Due to its smaller size, the Y chromosome-specific RH-PCR network showed reduced but still significant overlap with STRING (optimum FDR = 6.8 × 10^−12^, OR = 2.6).

### Figures of Merit for Genetic Interaction Networks

Receiver operating characteristic (ROC) curves were used to evaluate how well the essential RH-PCR and CRISPRi networks identify interactions present in STRING (Fig. 4*A*). Although areas under the curve (AUCs) exceeded random performance (0.5) only modestly, all differences were statistically significant (Z = 74.9, *P* < 2.2 × 10^−308^). Consistent with Fisher’s Exact Test results (Fig. 3*B*), the K562 network demonstrated superior predictive power compared to Jurkat, which in turn outperformed the essential RH-PCR network.

**Figure 4.**
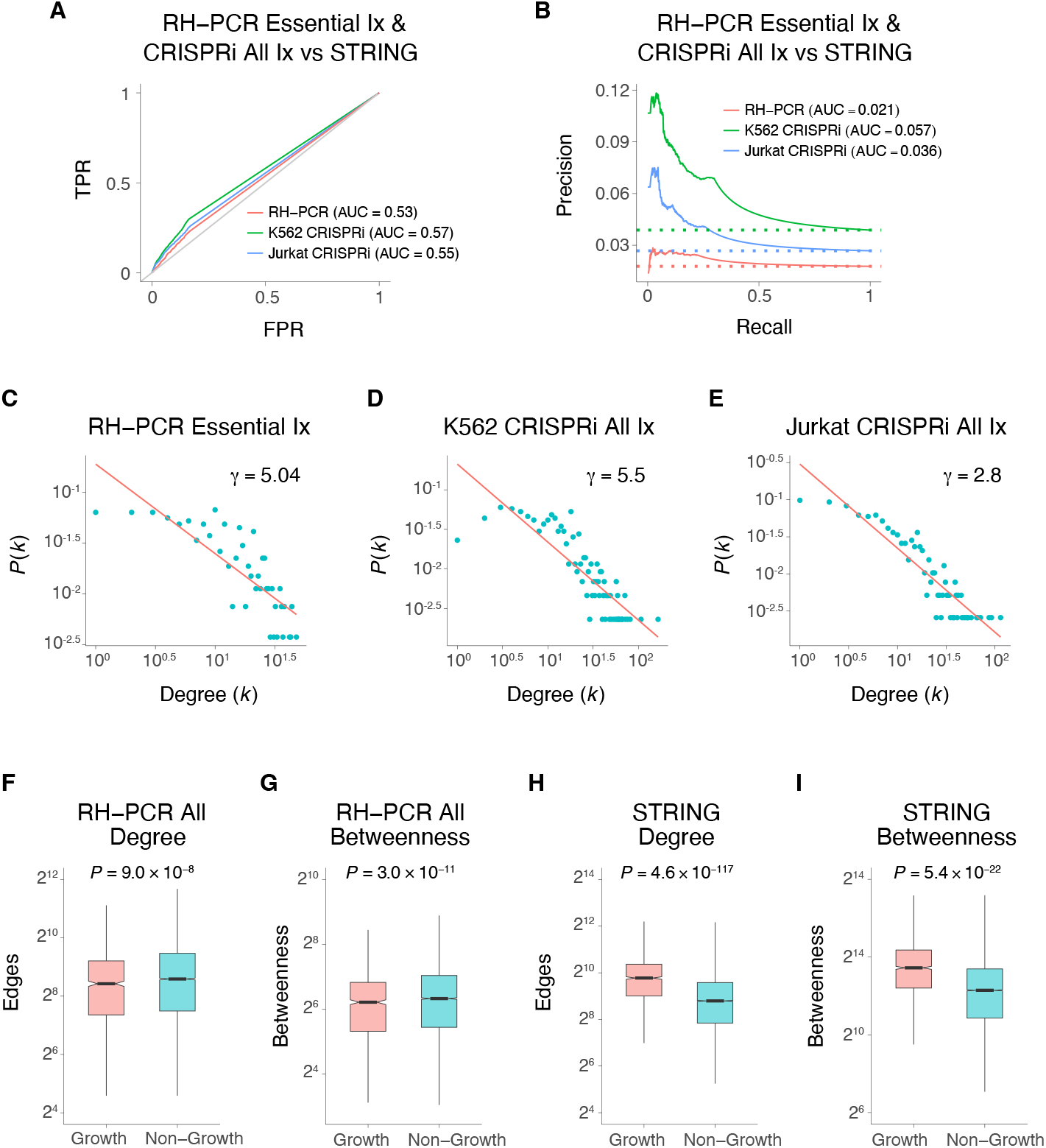
RH-PCR, CRISPRi and STRING interaction networks. *A*: ROC curves. Essential RH-PCR and combined buffering and synergistic K562 and Jurkat networks. TPR, true positive rate; FPR, false positive rate. Gray line, random. *B*: Precision recall curves. Dotted lines, random. *C*: Degree distribution for RH-PCR essential network. *D*: Degree distribution for K562 combined buffering and synergistic CRISPRi network. *E*: Degree distribution for combined Jurkat network. *F*: Degree centrality for protein coding RH-PCR network. *G*: Betweenness centrality for protein coding RH-PCR network. *H*: Degree centrality for STRING. *I*: Betweenness centrality for STRING. Growth, genes required for growth. Non-growth, not required.

When events of interest (significant genetic interactions), are greatly outnumbered by null events (no interactions) the precision recall (PRC) plot is superior to the ROC curve (41). PRC plots comparing the essential RH-PCR and CRISPRi networks with STRING showed results consistent with the ROC curves and Fisher’s Exact Test (Fig. 4*B*). All three networks showed modest power to identify STRING interactions, though each performed significantly better than random (W > 8.2 × 10^7^, *P* < 3.8 × 10^−12^, Wilcoxon Rank Sum Test).

### Topology of RH-PCR and CRISPRi Networks

Both the essential RH-PCR and CRISPRi networks were approximately scale-free (Fig. 4*C-E*), a frequent property of biological networks (42, 43). Scale-free networks follow a power law distribution, *P*(*k*) ∼ *k*^−*γ*^, where *P*(*k*) is the fraction of nodes having *k* connections, and *γ* is a scale factor. The RH-PCR and K562 networks had *γ*∼5, while the Jurkat network had a smaller *γ*∼2.8.

Whether degree and betweenness centralities in biological networks correlate positively with gene essentiality remains controversial (44, 45). To investigate these relationships in the protein coding RH-PCR and STRING networks, we obtained gene essentiality data from a CRISPR single gene knockout dataset (null alleles) in K562 cells (34).

Growth genes showed significantly reduced degree and betweenness centralities in the protein coding RH-PCR network (degree: *t*[1,1341] = 5.4, *P* = 9.0 × 10^−8^; betweenness: *t*[1,1533] = 6.7, *P* = 3.0 × 10^−11^; Welch Two Sample t-test) (Fig. 4*F* and *G*). Conversely, growth genes exhibited elevated degree and betweenness centralities in STRING (degree: *t*[1,1770] = 24.8, *P* = 4.6 × 10^−117^; betweenness: *t*[1,1692] = 9.8, *P* = 5.4 × 10^−22^; Welch Two Sample t-test) (Fig. 4*H* and *I*).

Growth suppressing genes (which increased growth when knocked out) showed reversed relationships in both the RH-PCR and STRING networks (Supplemental Fig. S7). Thus, growth genes tend to be weakly connected in genetic interaction networks, while growth suppressing genes are highly connected. This relationship is inverted in protein interaction networks.

Because the CRISPRi network was restricted to essential genes, it was not possible to explore the relationship between centrality and essentiality in this network.

### Gene Expression and RH-PCR and CRISPRi Networks

Microarray data from a human RH panel (33) was used to examine whether highly expressed genes are more likely to participate in genetic interactions. Both the essential and protein coding RH-PCR networks showed weak but significant positive correlations between genetic interaction strength (−log_10_*P*) and gene expression (essential: *R* = 0.06, *t*[1,1649] = 2.5, *P* = 0.013; full: *R* = 0.02, *t*[1,2320329] = 25.1, *P* = 3.3 × 10^−139^).

Similarly, RNA-Seq data showed significant positive correlations between expression and CRISPRi GI score in K562 and Jurkat cells, though the correlations were small (*R* = 0.03, *t*[1,85076] = 8.7, *P* = 4.6 × 10^−18^; *R* = 0.03, *t*[1,81001] = 9.9, *P* = 6.9 × 10^−23^, respectively) (31, 32).

### Genes Involved in RH-PCR and CRISPRi Interactions

The essential RH-PCR interaction network showed no significant overlap with the K562 or Jurkat CRISPRi networks (K562 combined interactions: optimum FDR = 1.0, OR = 0.21) (Supplemental Table S3). However, the genes participating in interactions showed highly significant overlap (K562 combined interactions: optimum FDR = 1.3 × 10^−5^, OR = 1.8) (Fig. 3*F*, Supplemental Table S4, Supplemental Figs. S8, S9).

Random gene samples matching the size of the essential RH-PCR network showed no significant overlap with CRISPRi networks at the optimal overlap parameters of actual data (K562 combined interactions: most significant FDR = 0.4; Jurkat combined interactions: FDR = 0.9; 2,000 bootstrap samples). This observation aligns with null expectations from Fisher’s Exact Test.

Consistent with the non-significant overlap of interactions, only 67 were shared between the 3,885 combined buffering and synergistic K562 CRISPRi interactions and 1,778 essential RH-PCR interactions (Fig. 5*A*, Supplemental Fig. S10*A*). In contrast, among the 446 and 279 genes involved in K562 CRISPRi and essential RH-PCR interactions respectively, 277 were shared (Fig. 5*B*, Supplemental Fig. S10*B*). Thus, the RH-PCR and CRISPRi approaches identify nearly identical sets of interacting genes while highlighting different interaction pairs.

**Figure 5.**
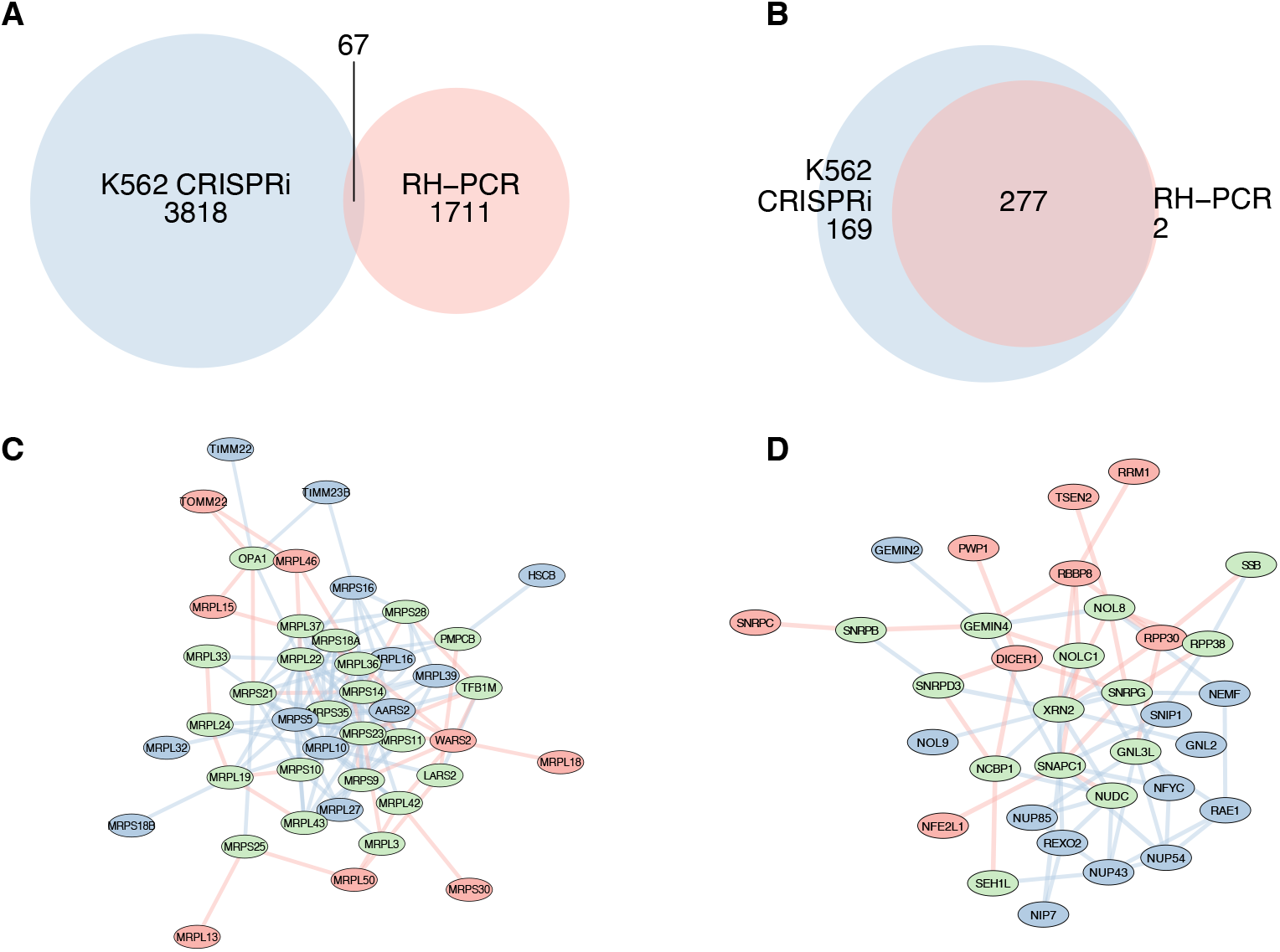
Overlaps of interactions and genes. *A*: Overlap of significant interactions in the essential RH-PCR and K562 combined buffering and synergistic CRISPRi networks. *B*: Overlap of genes involved in significant interactions in the two networks. *C*: Interactions between mitochondrial genes in amalgamated essential RH-PCR and K562 buffering and synergistic CRISPRi networks. Red nodes and edges, RH-PCR network only; blue nodes and edges, CRISPRi only; green nodes, both RH-PCR and CRISPRi. Edge width proportional to interaction strength. *D*: Interactions between genes that encode nuclear-and nucleolus-related proteins in amalgamated essential RH-PCR and K562 buffering and synergistic CRISPRi networks.

Examples of complementary genetic interactions in the essential RH-PCR and K562 combined CRISPRi networks are shown for mitchondrial and nuclear-related genes in Fig. 5*C* and *D*.

The mitochondrial interaction network obtained by combining the essential RH-PCR and buffering and synergistic K562 CRISPRi datasets involved 43 genes, of which 23 were shared (Fig. 5*C*). However, among 127 interactions in this network, only 4 were common to both approaches. Similarly, the network of essential nuclear- and nucleolus-related genes contained 34 genes, of which 14 were shared between RH-PCR and K562 CRISPRi networks, yet among 65 interactions only one was common (Fig. 5*D*).

### Functional Clustering of Essential Genes in the RH-PCR Network

To further explore the essential RH-PCR network, unsupervised clustering of genes based on their interaction −log_10_*P* values was performed (Fig. 6*A*). Clusters were then assigned to gene ontology (GO) enrichment categories (FDR < 10^−5^) (36).

**Figure 6.**
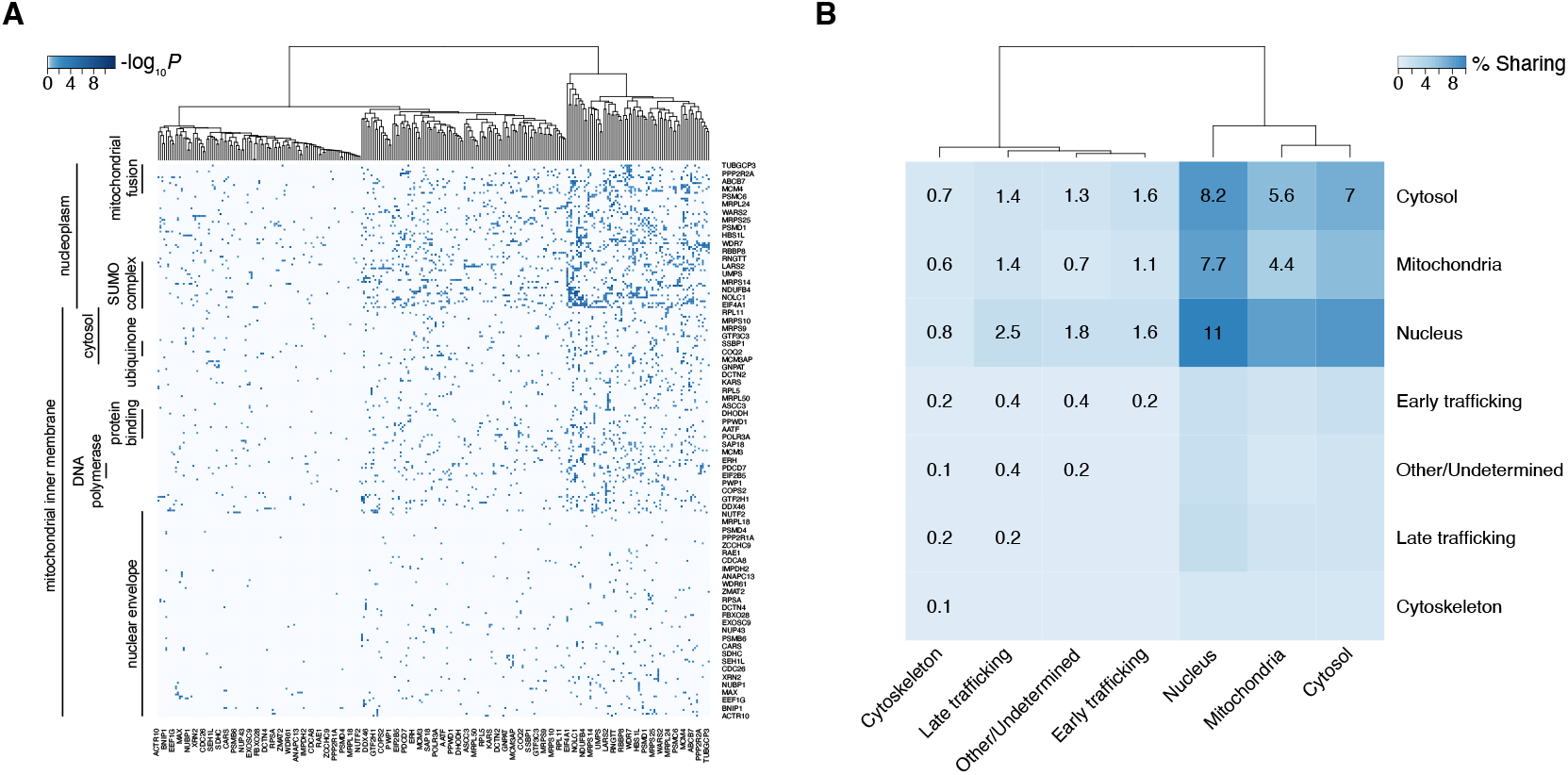
Functional clustering of essential RH-PCR network. *A*: Gene clusters based on −log_10_*P* interaction values. Functional enrichments use gene ontology (GO) categories. For reasons of space, only 13% of the genes in the cluster diagram are labeled. *B*: Clustering of shared cellular compartments based on percent interacting genes expressed in each compartment. Percent sharing indicated by color and number in each heatmap square. Dendrograms resulting from unsupervised clustering shown above heatmaps.

GO terms related to mitochondria (mitochondrial inner membrane, mitochondrial fusion), nucleus (nucleoplasm, nuclear envelope), and cytosol were predominant. For example, among 241 nuclearly-encoded mitochondrial genes, 211 were in the mitochondrial inner membrane cluster. Similarly, among 308 genes with nucleus-related functions, 105 clustered within the nuclear envelope category.

CRISPRi interaction network clustering yielded similar patterns, with prominent enrichment of nucleus/cytoplasm, cytosol, and mitochondria (7). However, some GO terms in the CRISPRi network, such as mRNA stability and splicing, did not appear in the essential RH-PCR network.

### Cellular Compartments of Essential genes in the RH-PCR Network

Genetic interactions are reflected in the shared cellular compartments of corresponding proteins. To better understand this aspect of the essential RH-PCR network, unsupervised clustering of cellular compartments was performed using the proportion of interacting genes expressed in each compartment (Fig. 6*B*).

Reminiscent of the GO enrichment clusters, the three compartments with highest proportions of shared interactions were nucleus, cytosol, and mitochondria. The essential RH-PCR network showed both similarities and differences with the CRISPRi network, which exhibited strong clustering of mitochondria for buffering interactions and other/undetermined plus early trafficking compartments for synergistic interactions (7).

### Genetic Interaction Networks and GWAS

Genome-wide association studies (GWASs) identify individual variants contributing to complex disorders. However, these variants reflect underlying genetic interaction networks that are difficult to dissect due to limited statistical power (46, 47). The potential of RH-PCR and CRISPRi networks to elucidate these interactions was investigated by constructing a network linking genes associated with the same phenotype in a dataset of GWAS results (18).

The essential RH-PCR network showed significant overlap with the GWAS network (optimum FDR = 1.9 × 10^−14^, OR = 2.0), while the K562 and Jurkat CRISPRi networks did not (Fig. 7*A*, Supplemental Tables S5, S6). The GWAS network also overlapped significantly with the protein coding RH-PCR network (optimum FDR = 7.2 × 10^−319^, OR = 1.7 ± 0.1 s.d.) and a pruned network consisting of essential genes and genes one link away (optimum FDR = 4.5 × 10^−250^, OR = 16.1 ± 2.6 s.d.) (Fig. 7*B*, Supplemental Table S5).

**Figure 7.**
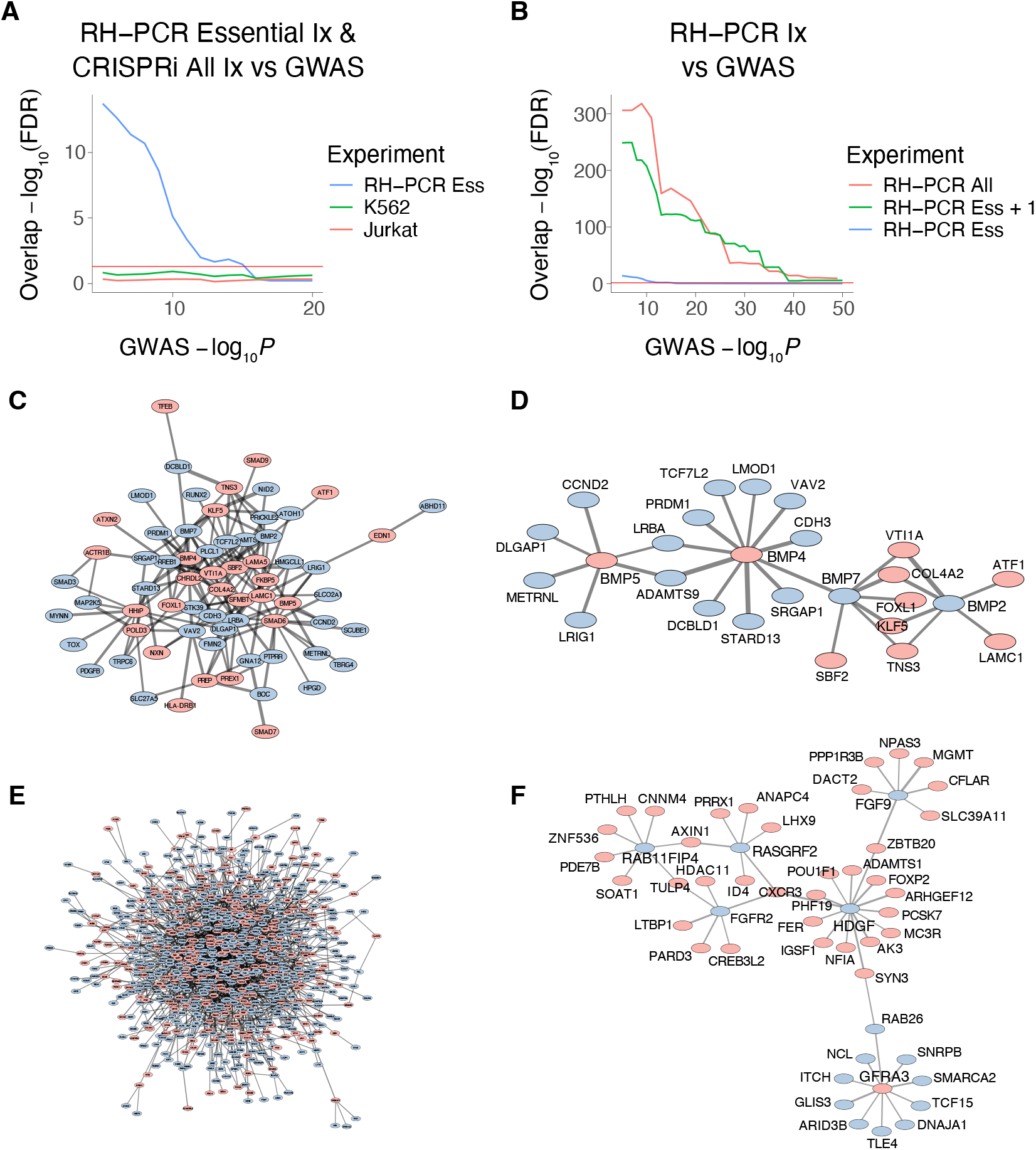
RH-PCR, CRISPRi and GWAS networks. *A*: Overlaps of essential RH-PCR and K562 and Jurkat combined buffering and synergistic CRISPRi networks with GWAS network. Horizontal red line, FDR (false discovery rate) = 0.05. *B*: Overlaps of protein coding RH-PCR, essential RH-PCR plus genes one link away, and essential RH-PCR networks with GWAS network. *C*: Colorectal cancer GWAS genes from lower power study (blue nodes) and additional genes in the higher power study (pink) predicted by the RH-PCR network. Edge width reflects RH-PCR −log_10_*P. D*: Subnetwork for colorectal cancer featuring BMPs. *E*: Height GWAS. *F*: Subnetwork for height featuring growth-related genes.

GWAS resolution is not uniformly single gene and the GWAS network was constructed using both single- and multigene loci to avoid overfitting. To evaluate whether GWAS network overlaps were substantially confounded by linkage, we constructed an additional GWAS network using only single gene entries.

The single gene GWAS network showed significant overlap with the protein coding RH-PCR network, but with an increased OR (optimum FDR = 2.2 × 10^−308^, OR = 2.9 ± 0.1 s.d.) compared to the GWAS network including multigene entries. The optimum overlap between the CRISPRi and single gene GWAS networks remained insignificant. These observations suggest tlinkage does not substantially contribute to GWAS network overlaps.

The significant RH-PCR and GWAS network overlaps may reflect the milder effect of extra gene copies in RH cells compared to loss-of-function CRISPRi alleles. Most common GWAS alleles affect gene regulation and have weaker effects than loss-of-function variants (48).

Significant overlap was found between downsampled protein coding RH-PCR and GWAS networks (single- and multigene entries) in which at least one gene per interaction was on the X chromosome (optimum FDR = 4.7 × 10^−10^, OR = 2.6) (Supplemental Table S5). However, overlap significance was weaker than for the complete RH-PCR and GWAS networks, unlike the overlap between X chromosome-specific RH-PCR and STRING networks (Fig. 3*E*).

GWASs that include sex chromosomes are scarce (49). Consistent with this lack, X chromosome loci in the GWAS dataset accounted for only 0.5% of all loci. This ascertainment bias may explain the weaker (though still significant) overlap of X chromosome-specific RH-PCR and GWAS networks.

For sufficiently large GWASs, significant overlaps existed between the protein coding RH-PCR network and subnetworks for individual traits, including body mass index (*P* = 9.7 × 10^−154^, OR = 1.9), colorectal cancer (*P* = 1.0 × 10^−32^, OR = 1.4), and height (*P* = 2.2 × 10^−308^, OR = 1.4 ± 0.04 s.d.) (Supplemental Table S7).

The overlaps between the protein coding RH-PCR and GWAS trait-specific subnetworks prompted an examination of whether the RH-PCR network could predict loci in higher-powered GWASs using lower-powered GWASs. Two colorectal cancer GWASs were used. The earlier, smaller GWAS had ∼35,000 cases and ∼71,000 controls, identifying 70 loci (50). The later GWAS had greater power, with ∼100,000 cases and ∼155,000 controls, identifying 162 loci (51). A total of 55 loci were shared between the two studies.

RH-PCR network interactions linking genes from the smaller GWAS to genes in the larger study by a single step showed significant enrichment compared to background (OR = 1.5, *P* = 6.2 × 10^−6^). Of 107 disease-related genes (DRGs) for colorectal cancer found in the larger but not smaller GWAS, 40 were linked by the RH-PCR network to the smaller study (Fig. 7*C*, Supplemental Table S8).

A subnetwork of the two RH-PCR linked networks featuring bone morphogenetic proteins (BMPs), which play an important role in colorectal cancer, is shown in Fig. 7*D* (52). Genes in the smaller study connected by the RH-PCR network to predicted DRGs in the larger study included those with a role in oncogenic proliferation, such as *CCND2* linked to *BMP5, VAV2* linked to *BMP4, BMP7* linked to *KLF5*, and *BMP2* linked to *LAMC1*.

In another example, two height GWASs were evaluated. The lower-powered study examined ∼628,000 individuals and identified 893 loci, while the higher-powered study examined 5.4 million individuals and identified 2,483 loci (53, 54). A total of 529 loci were shared between the two studies. RH-PCR network interactions linking genes from the smaller GWAS to genes in the larger showed significant enrichment (OR = 1.8, *P* = 2.2 × 10^−308^). Of 1,954 genes specific to the higher-powered GWAS, 895 were linked by the RH-PCR network to genes in the lower-powered study (Fig. 7*E*, Supplemental Table S8).

A subnetwork of the two RH-PCR linked networks for height emphasizing proliferation-related genes is shown in Fig. 7*F* (54). Examples of genes from the smaller study connected by the RH-PCR network to the larger study included those involved in cell growth, such as *FGFR2* linked to *LTBP1, FGF9* linked to *DACT2*, and *SMARCA2* linked to *GFRA3*.

The RH-PCR network may thus identify novel candidate genes using data from modestly powered GWASs, such as for uncommon or difficult-to-ascertain phenotypes.

## DISCUSSION

Charting biological networks from multiple perspectives provides deeper insights into cellular mechanisms. While network comparisons have been feasible in lower eukaryotes such as yeast (55), such studies have been challenging in mammalian cells until recently. Understanding the differences between interaction networks will require detailed, comprehensive investigations.

This study is the first to examine independently constructed genetic networks in mammalian cells. The RH-PCR network employed extra gene copies while the CRISPRi network used partial loss-of-function alleles. Comparisons of the networks revealed: Both CRISPRi and RH-PCR networks overlap with proteinprotein interaction networks. (2) RH-PCR and CRISPRi networks identify a core set of interacting genes despite differences in actual interacting gene pairs. (3) The RH-PCR network overlaps more strongly with GWAS networks than CRISPRi, illuminating the relationship between variant types and complex genetic traits. (4) Disease-related genes can be predicted using RH-PCR networks.

The RH-PCR network was constructed using panels with donor DNA from four species. Although random donor DNA fragments provide the genetic perturbations in RH cells, disparities between RH-PCR and CRISPRi networks are unlikely to result from donor DNA identity. This conclusion is supported by strong overlaps among genetic networks derived from RH panels with different donor species (4).

The RH recipient cells were hamster fibroblasts or epithelial cells, while the K562 and Jurkat cells used in the CRISPRi study were human leukemia cells. Since genetic mechanisms of cell growth are conserved, differences between RH-PCR and CRISPRi networks more likely reflect their distinct variant types (gain- and loss-of-function respectively) rather than dissimilar cell types (56).

Although experimental verification of selected RH-PCR network interactions will be valuable, given its size these studies can only explore a small corner of the network. Validation of a large network is best done at a genome-wide level. The highly significant overlap between the RH-PCR network and multiple interaction networks suggests the RH approach can inform diverse biological perspectives.

The relatively non-toxic effects of increased gene copy number in the RH approach is reflected in the uniform retention of donor DNA, which allows mapping of interactions for the entire genome (1, 4, 16, 29, 39, 33, 57, 58). Conversely, the genes in the CRISPRi study were chosen for moderate growth phenotypes, since interactions cannot be assessed when gene loss causes cell death. Further, the relatively low CRISPRi throughput renders evaluation of all interactions prohibitive.

Even though the essential RH-PCR network was downsampled to match CRISPRi genes, restricting the CRISPRi network to genes with mild growth effects may introduce selection bias. Additional insights may emerge with networks using higher copy numbers or stronger loss-of-function alleles.

Both CRISPRi and protein coding RH-PCR networks showed significant overlap with the STRING protein-protein interaction repository. The RH-PCR network showed superior overlap, likely due to its larger size. However, the CRISPRi interaction map showed greater overlap with STRING than the essential RH-PCR network.

Despite significant STRING overlaps, precision-recall curves demonstrated limited power of RH-PCR and CRISPRi networks to predict protein interactions. This finding is not surprising, as genes may interact indirectly or through direct protein association.

No significant overlap existed between RH-PCR and CRISPRi interaction networks, reflecting their different mechanisms. This finding aligns with studies showing that growth genes identified by partial inactivation using CRISPRi differ from those identified by CRISPRa (8).

Conversely, CRISPRi and RH-PCR networks showed significant overlap of participating genes rather than interactions. Thus, partial loss-of-function and extra gene copies implicate similar genes in biological processes while highlighting distinct interaction partners. These similarities and differences may reflect hidden commonalities that become manifest when evaluating higher-order interactions (59).

Genome-wide mapping of three-way interactions is difficult using CRISPR, but feasible with RH technology because of the large number of extra genes in each cell. Nevertheless, even in RH cells, three-way interactions occur at lower frequency than two-way (∼cube of the retention probability vs square). Large datasets will be required for adequate statistical power.

The marked preponderance of attractive over repulsive interactions in the RH-PCR network contrasts with the CRISPRi network. The CRISPRi network has roughly equal abundances of buffering and synergistic interactions, each showing similar overlaps with the unsigned STRING network. Direct equivalence of the two RH-PCR interaction types and two CRISPRi interaction types is unlikely.

The protein coding RH-PCR network showed diminished interaction strength for X chromosome genes compared to autosomes. Weaker interactions of X chromosome genes is consistent with their blunted expression increases in response to copy number increments (33, 39), perhaps reflecting adaptation to sex-based copy number changes (40). The attenuated expression responses and weaker genetic interactions of X chromosome genes may constitute a dosage compensation mechanism that operates alongside classical X inactivation (33, 39).

Although constructed differently, the essential RH-PCR and CRISPRi networks both exhibited scale-free topology. Biological networks typically have scale factor *γ* ∼2–3 (42, 43). Such networks have few abundantly connected “hub” nodes and are robust to node removal or replacement. Networks with larger *γ* values, ∼5, have even fewer highly connected hubs and greater robustness to random node removal. The RH-PCR and K562 networks both had *γ*∼5, while the Jurkat network had a smaller *γ*∼2.8.

Growth promoting and suppressing genes showed decreased and increased centrality, respectively, in the protein coding RH-PCR network. This observation implies that highly connected nodes are likely to inhibit cell proliferation in genetic interaction networks. Surprisingly, the relationship between growth and centrality was reversed in STRING, exemplifying the encrypted relationship between genetic and physical interactions.

Functional clustering of the essential RH-PCR and CRISPRi networks showed similar enrichments, consistent with their shared node genes. GO analysis highlighted mitochondria and nucleus, while compartment classifications emphasized nucleus, cytoplasm, and mitochondria. Other compartment assignments present in the CRISPRi network, such as early trafficking and cytoskeleton, were not clearly delineated in the RH-PCR network.

The RH-PCR network showed significant overlap with a network constructed from human GWAS data, while the CRISPRi network did not. The lack of CRISPRi and GWAS network overlap was not due to insufficient power, since an RH-PCR network constructed from the same genes as CRISPRi showed significant overlap. Further, the protein coding RH-PCR network, but not the CRISPRi network, showed increased overlap with a single gene GWAS network compared to a GWAS network that included multigene entries. RH-PCR and GWAS network overlap is thus unlikely to be appreciably inflated by linkage, consistent with each network having near single gene resolution.

Overlaps of the RH-PCR and GWAS networks suggests that extra gene copies in the RH-PCR network have similar effects to common GWAS variants. This observation mirrors findings that growth genes identified using bulk segregant analysis of RH cells showed significant overlap with GWAS genes, while CRISPR-identified growth genes did not (16).

The RH approach uses natural gene promoters, while CRISPRi employs partial loss-of-function alleles. These technologies only produce phenotypes if targeted genes are normally transcriptionally active, consistent with weak but statistically significant correlations between interaction strength and gene expression in both networks.

Since CRISPRa overexpresses genes constitutively (8), RH-PCR and CRISPRi may provide more physiologically relevant genotype/phenotype information. Nevertheless, it will be informative to compare the RH-PCR and CRISPRi networks with more extensive genetic interaction maps constructed using CRISPRa when available. Understanding how genetic interactions contribute to cell growth will be deepened by developing genome-wide approaches that evaluate in depth the consequences of various loss- and gain-of-function alleles.

Recent developments may help elucidate mechanisms underlying genetic networks. These advances include massively parallel reporter assays (MPRA), which enable genome-scale dissection of promoter regions and transcriptional circuits, and Hi-C, which maps 3D chromatin folding and regulatory elements in the cell nucleus (60, 61).

## DATA AVAILABILITY

All data and code is available from https://doi.org/10.6084/m9. figshare.30311389.

## SUPPLEMENTAL MATERIAL

Supplemental Tables S1-S9 and Supplemental Figs. S1-S10: https://doi.org/10.6084/m9.figshare.30311551.

## ACKNOWLEDGMENTS

I thank Jake Lusis for helpful comments on the manuscript.

## AUTHOR CONTRIBUTIONS

D.J.S. conceived, designed and performed research.

## Supplemental material

**Figure S1:**
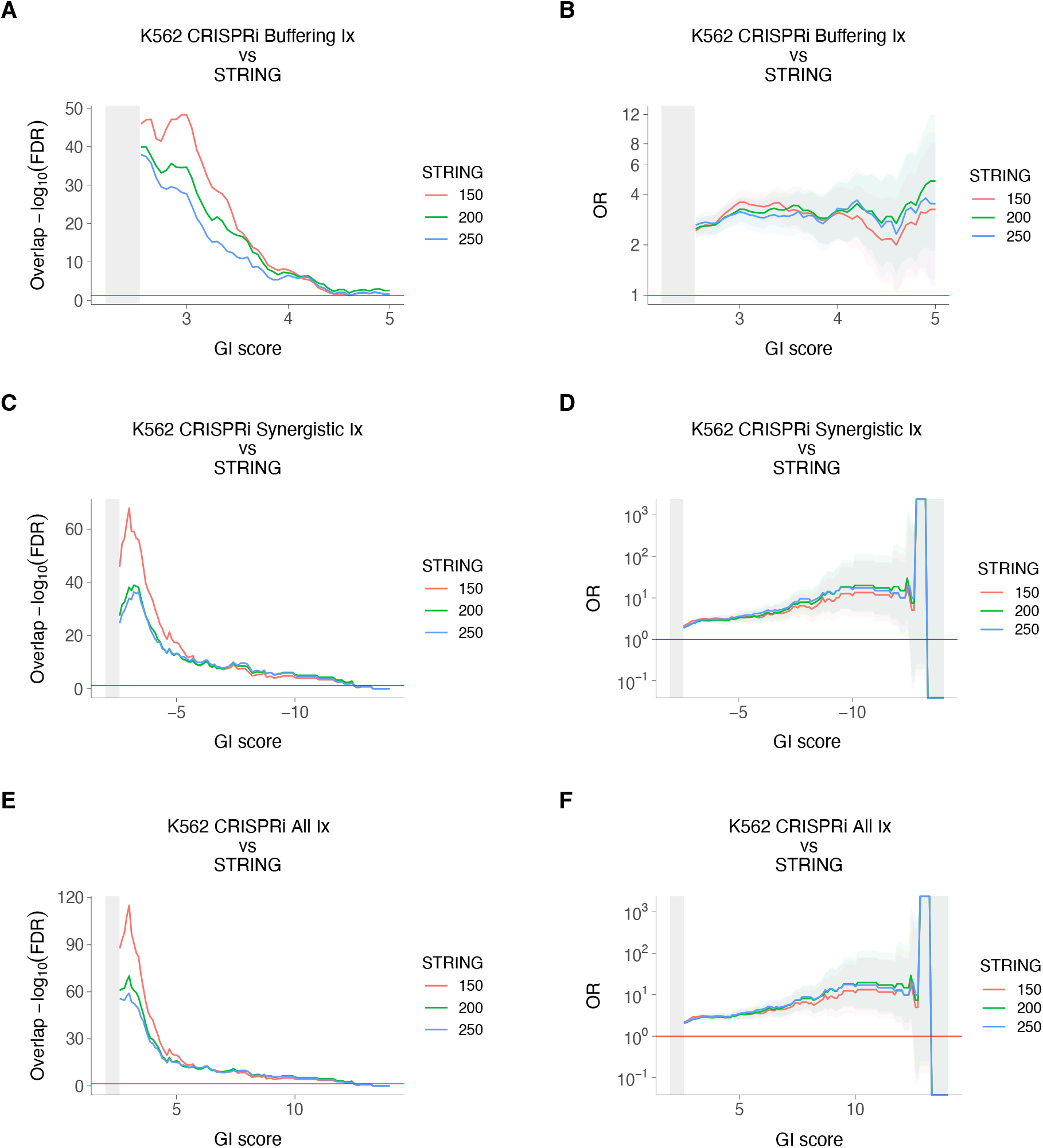
CRISPRi and STRING networks in K562 cells. *A*: Overlap of K562 buffering CRISPRi interactions (Ix) and STRING. Horizontal red line, FDR (false discovery rate) = 0.05. *B*: Odds ratios (ORs) for overlap of K562 buffering CRISPRi interactions and STRING. Confidence intervals shown (95%). Horizontal red line, OR = 1. *C*: Overlap for K562 synergistic CRISPRi interactions and STRING. *D*: ORs for K562 synergistic CRISPRi interactions and STRING. *E*: Overlap of K562 combined buffering and synergistic CRISPRi interactions and STRING. *F*: ORs for combined K562 CRISPRi interactions and STRING. FDR, false discovery rate. Gray rectangles at left of each graph indicate insignificant CRISPRi genetic interaction (GI) scores. ORs on a log_10_ scale.

**Figure S2:**
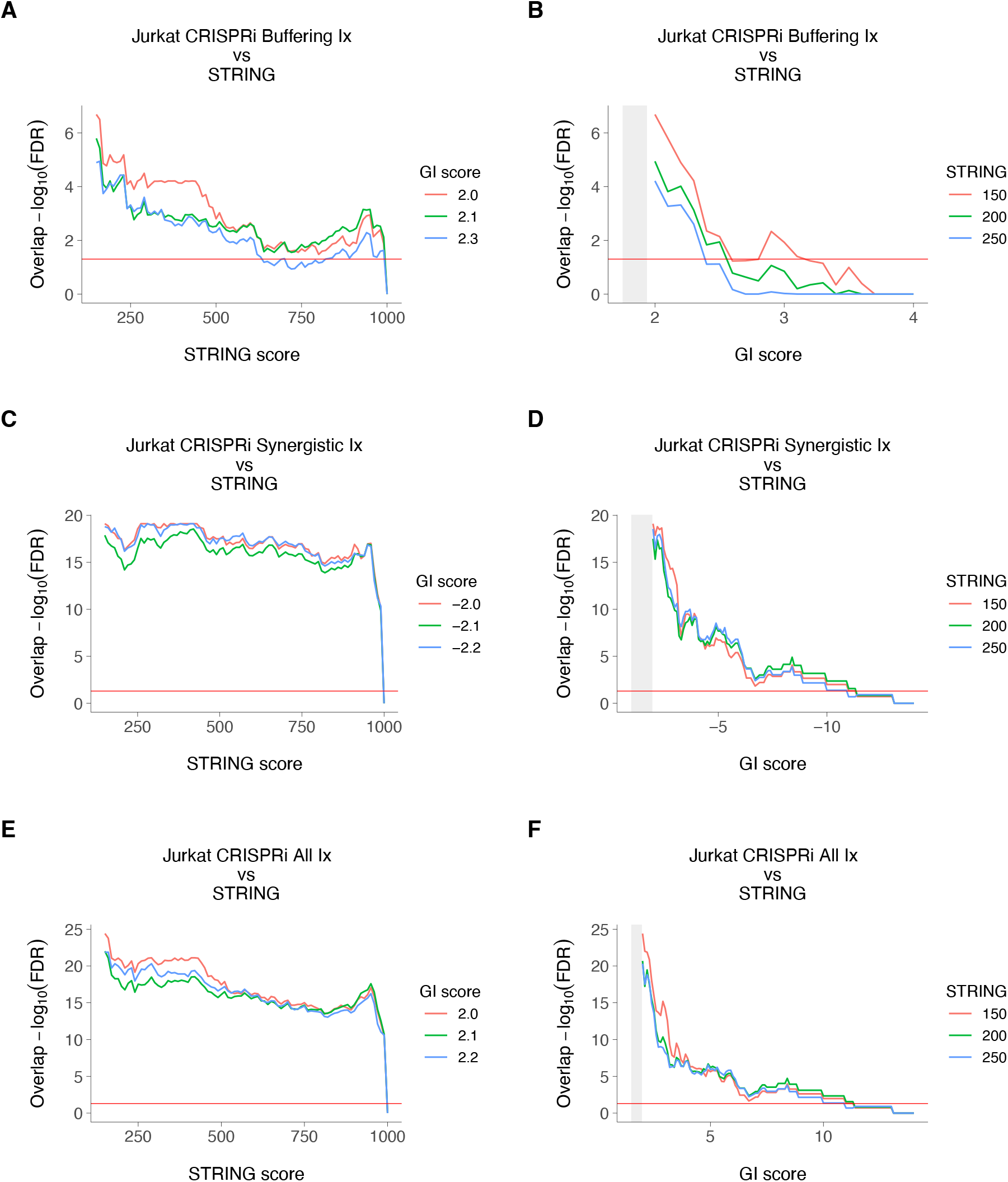
CRISPRi and STRING networks in Jurkat cells. *A,B*: Overlap of Jurkat buffering CRISPRi interactions and STRING. *C,D*: Overlap of Jurkat synergistic CRISPRi interactions and STRING. *E,F*: Overlap of Jurkat combined buffering and synergistic CRISPRi interactions and STRING. Gray rectangles at left of graphs in B, D and F indicate insignificant CRISPRi genetic interaction (GI) scores. Horizontal red lines, FDR (false discovery rate) = 0.05.

**Figure S3:**
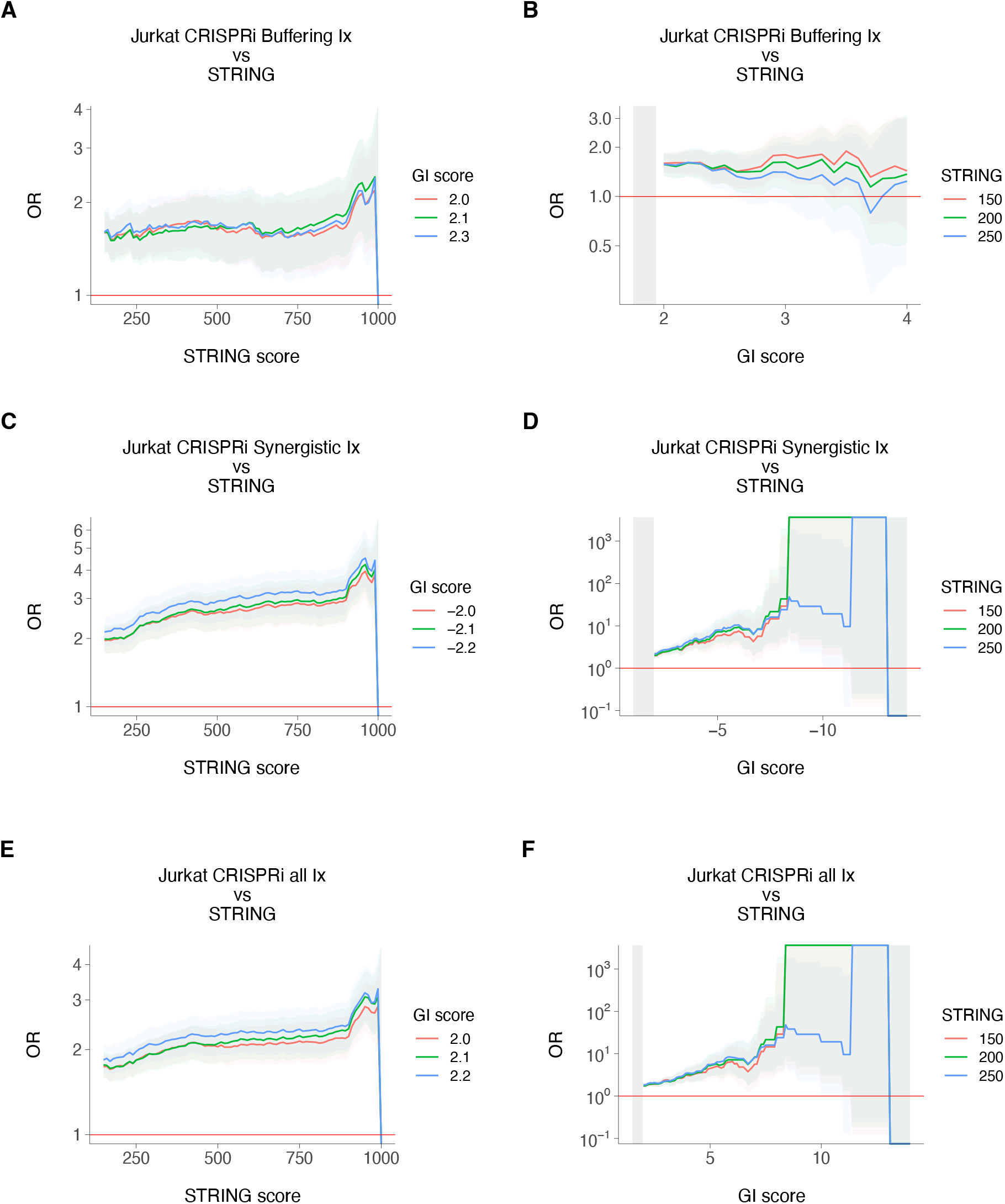
CRISPRi and STRING network odds ratios in Jurkat cells. *A,B*: Odds ratios (ORs) for overlap of Jurkat buffering CRISPRi interactions and STRING. *C,D*: ORs for overlap of Jurkat synergistic CRISPRi interactions and STRING. *E,F*: ORs for overlap of Jurkat combined buffering and synergistic CRISPRi interactions and STRING. Gray rectangles at left of graphs in B, D and F indicate insignificant CRISPRi genetic interaction (GI) scores. Horizontal red lines, OR = 1. ORs on a log_10_ scale.

**Figure S4:**
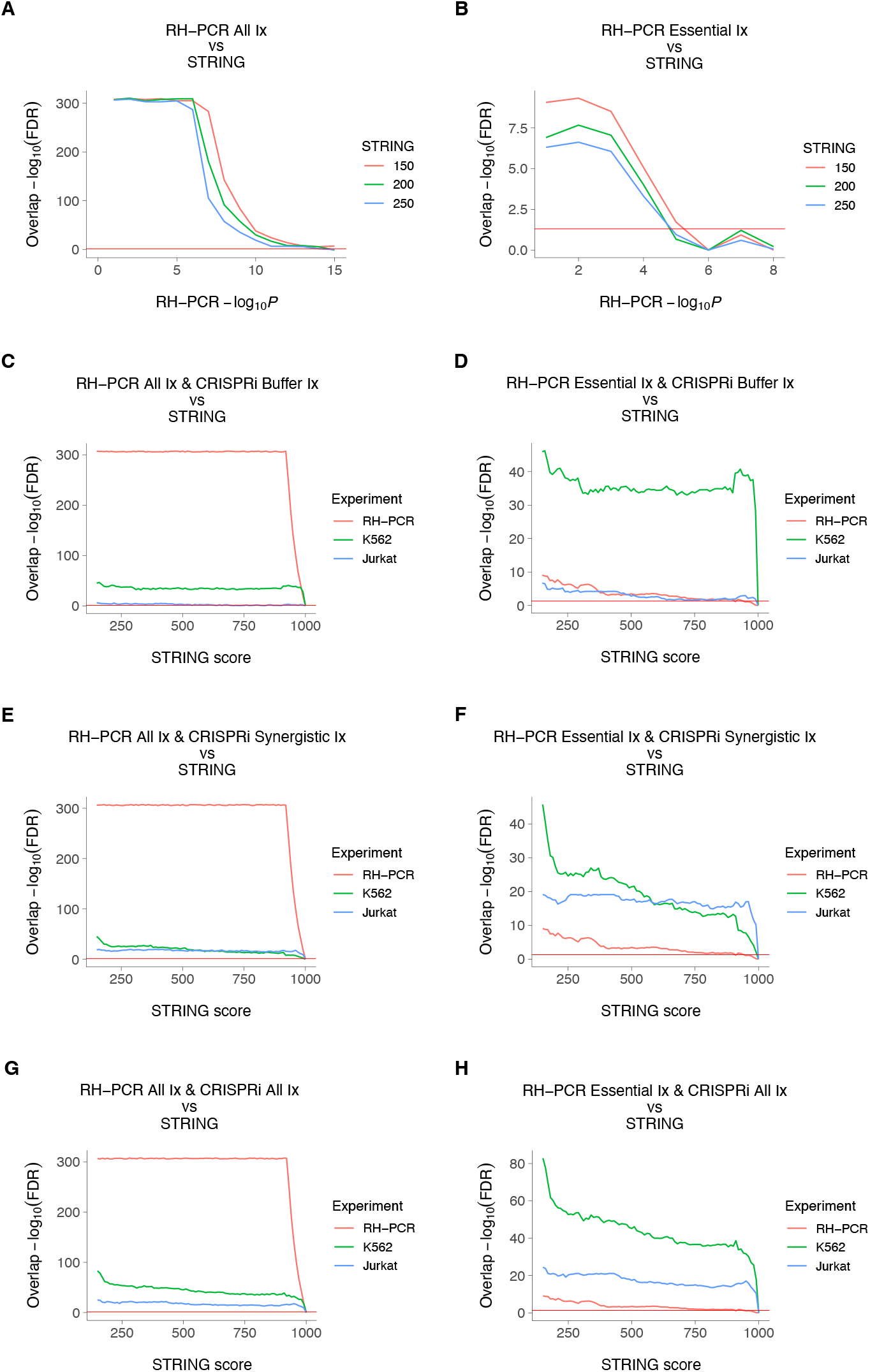
Overlaps of RH-PCR, CRISPRi and STRING networks. *A*: Overlap of protein coding RH-PCR network and STRING. *B*: Overlap of RH-PCR interactions restricted to CRISPRi essential genes and STRING. *C*: Overlap of protein coding RH-PCR, buffering CRISPRi, and STRING networks. *D*: Overlap of essential RH-PCR, buffering CRISPRi, and STRING networks. *E*: Overlap of protein coding RH-PCR, synergistic CRISPRi, and STRING networks. *F*: Overlap of essential RH-PCR, synergistic CRISPRi, and STRING networks. *G*: Overlap of protein coding RH-PCR, combined buffering and synergistic CRISPRi, and STRING networks. *H*: Overlap of essential RH-PCR, combined CRISPRi, and STRING networks. Horizontal red lines, FDR (false discovery rate) = 0.05.

**Figure S5:**
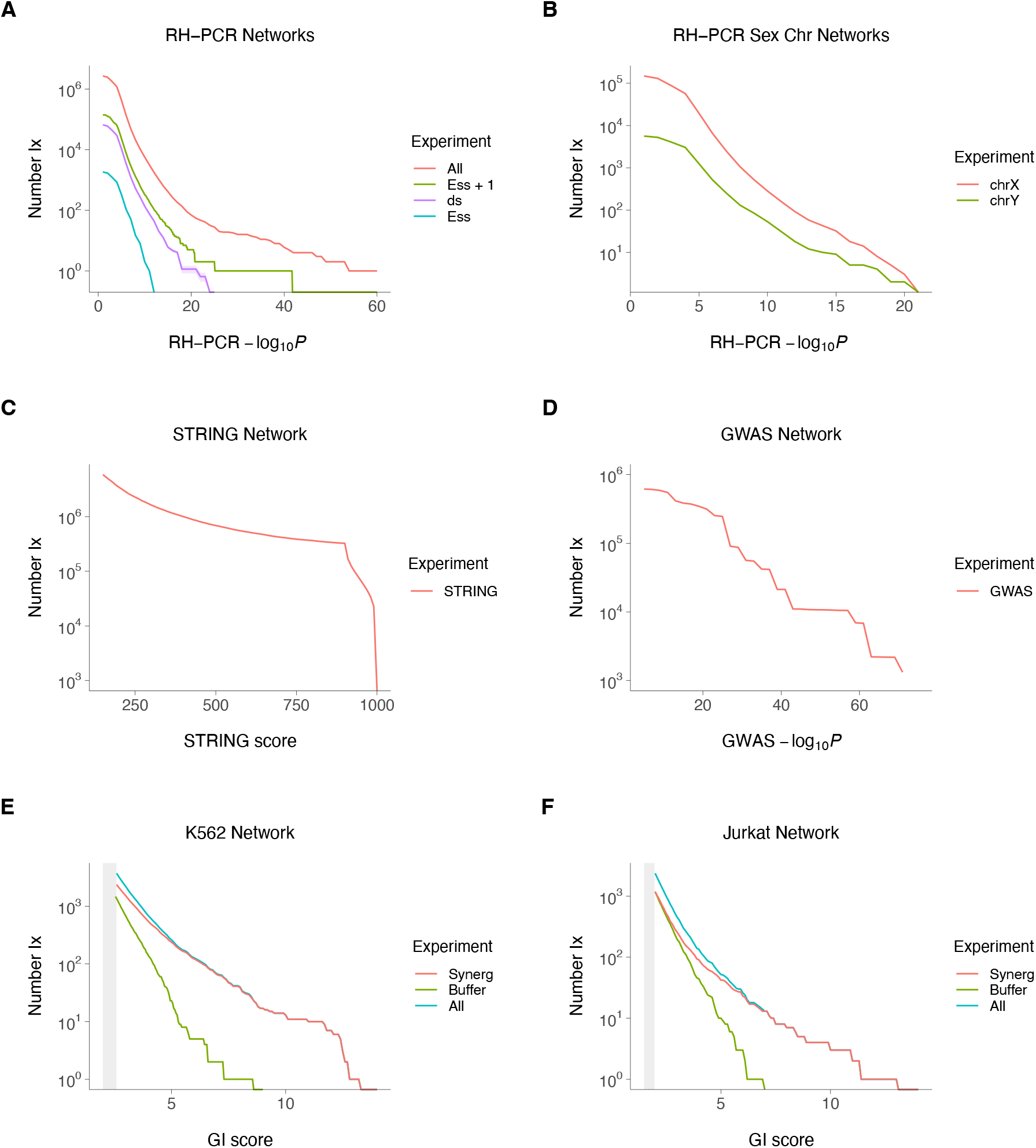
Network numbers. *A*: RH-PCR networks. All, protein-coding RH-PCR network. Ess + 1, essential RH-PCR network plus genes linked by one interaction. ds, downsampled RH-PCR network, 95% confidence limits shown. Ess, essential RH-PCR network. *B*: Protein coding RH-PCR network downsampled so that at least one gene in each interaction is on either the X or Y chromosome. *C*: STRING network. *D*: Geneome-wide association study (GWAS) network. *E*: K562 CRISPRi network. Synerg, synergistic interactions. Buffer, buffering interactions. All, combined synergistic and buffering interactions. *F*: Jurkat CRISPRi network. Gray rectangles at left of graphs in E and F indicate insignificant CRISPRi genetic interaction (GI) scores. Number Ix, number of interactions in each network as stringency increased.

**Figure S6:**
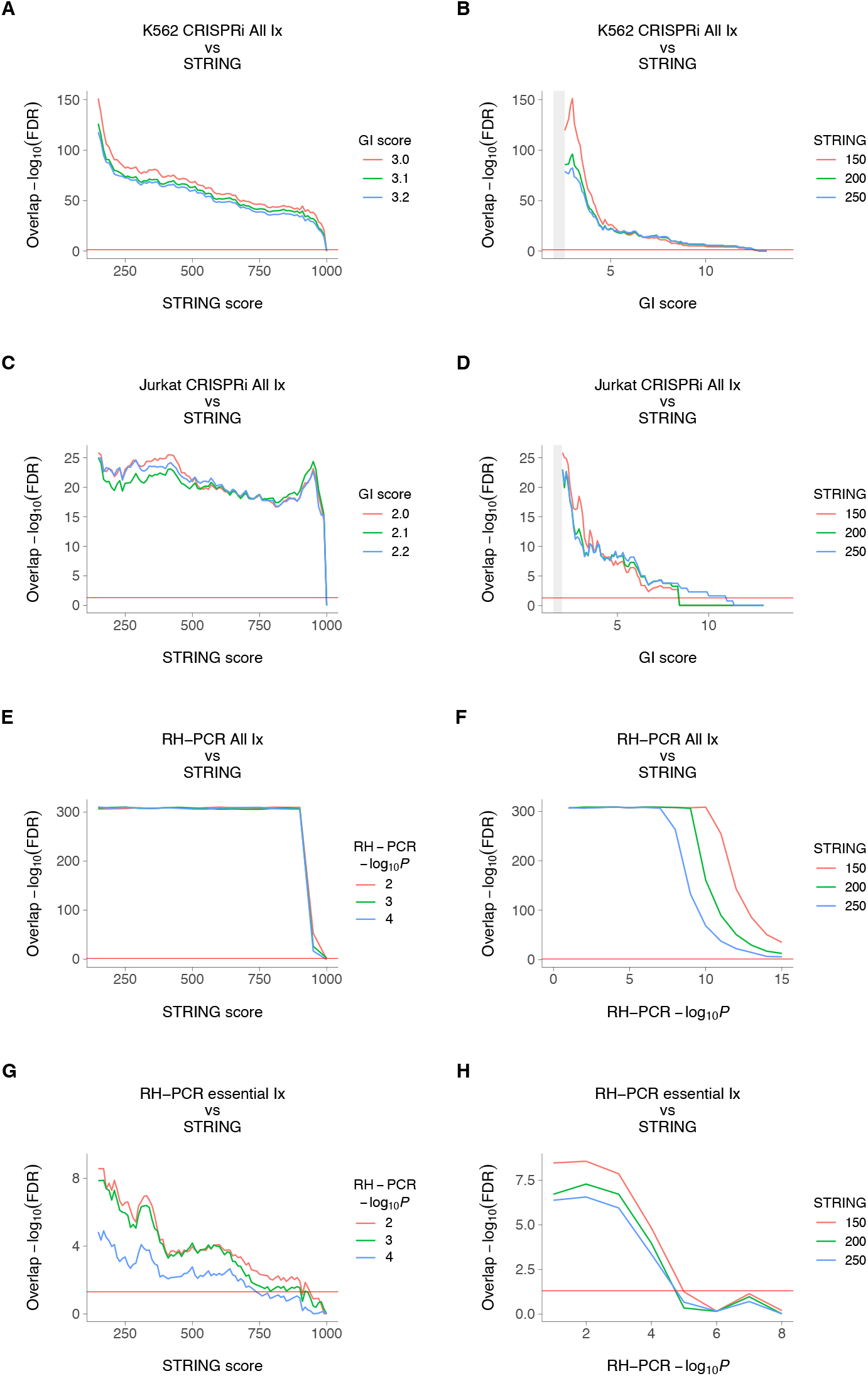
RH-PCR, CRISPRi and STRING interaction networks analyzed using logistic regression. *A,B*: Overlap of K562 combined buffering and synergistic CRISPRi interactions (All) and STRING. *C,D*: Overlap of Jurkat combined buffering and synergistic CRISPRi interactions and STRING. *E,F*: Overlap of protein coding RH-PCR network (All) and STRING. *G,H*: Overlap of RH-PCR interactions restricted to essential genes in the CRISPRi dataset and STRING. Gray rectangles at left of graphs in B and D indicate insignificant CRISPRi genetic interaction (GI) scores. Horizontal red lines, FDR (false discovery rate) = 0.05.

**Figure S7:**
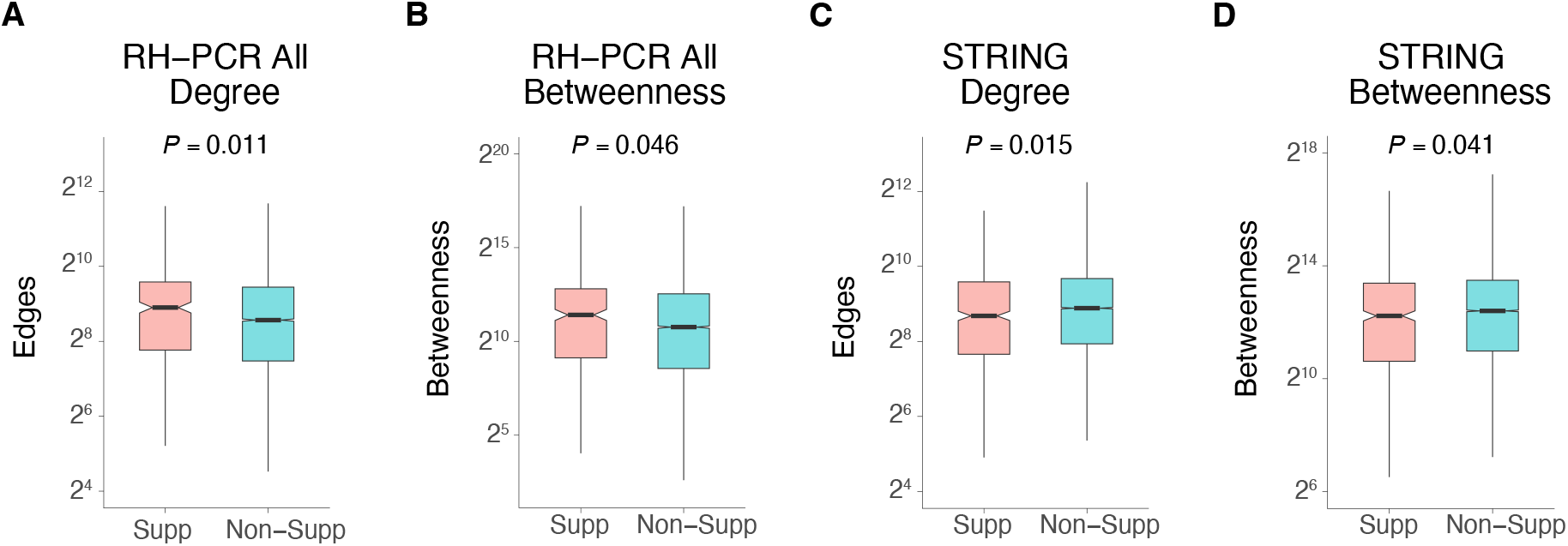
Centrality for growth suppressing genes in protein coding RH-PCR and STRING networks. *A*: Degree centrality for RH-PCR network, *t* [1,415] = 2.6, *P* =0.011. *B*: Betweenness centrality for RH-PCR network, *t* [1,408] = 2.0, *P* =0.046. *C*: Degree centrality for STRING network. *t* [1,649] = 2.5, *P* =0.015. *D*: Betweenness centrality for STRING network. *t* [1,711] = 2.0, *P* =0.041. Supp, genes that suppress growth. Non-Supp, non-suppressing. Welch Two Sample t-tests used.

**Figure S8:**
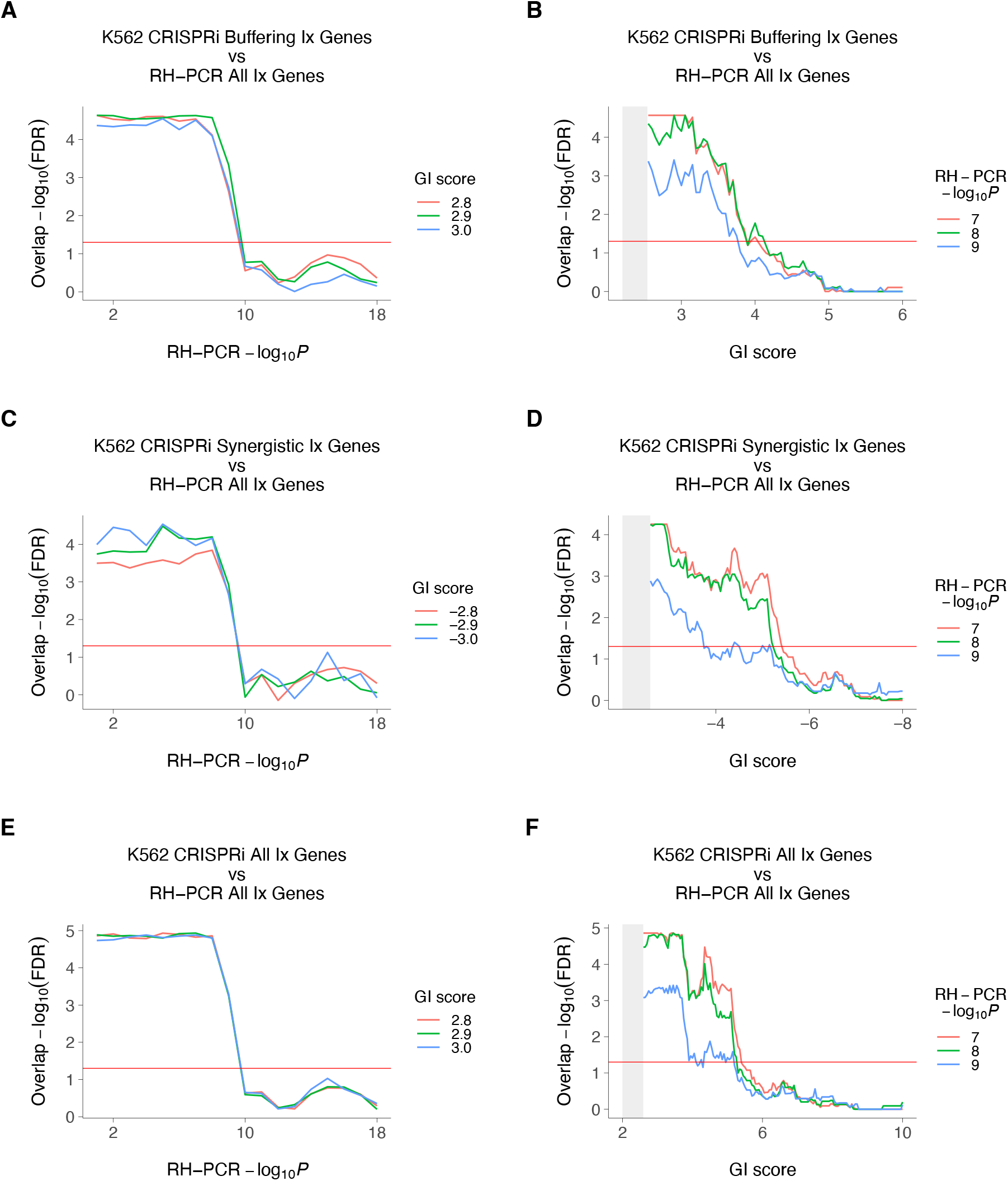
Genes in RH-PCR and K562 CRISPRi networks. *A,B*: Overlap of genes in K562 buffering CRISPRi and protein coding RH-PCR networks. *C,D*: Overlap of genes in K562 synergistic CRISPRi and protein coding RH-PCR networks. *E,F*: Overlap of genes in K562 combined buffering and synergistic CRISPRi and protein coding RH-PCR networks. Gray rectangles at left of graphs in B, D and F indicate insignificant CRISPRi genetic interaction (GI) scores. Horizontal red lines, FDR (false discovery rate) = 0.05.

**Figure S9:**
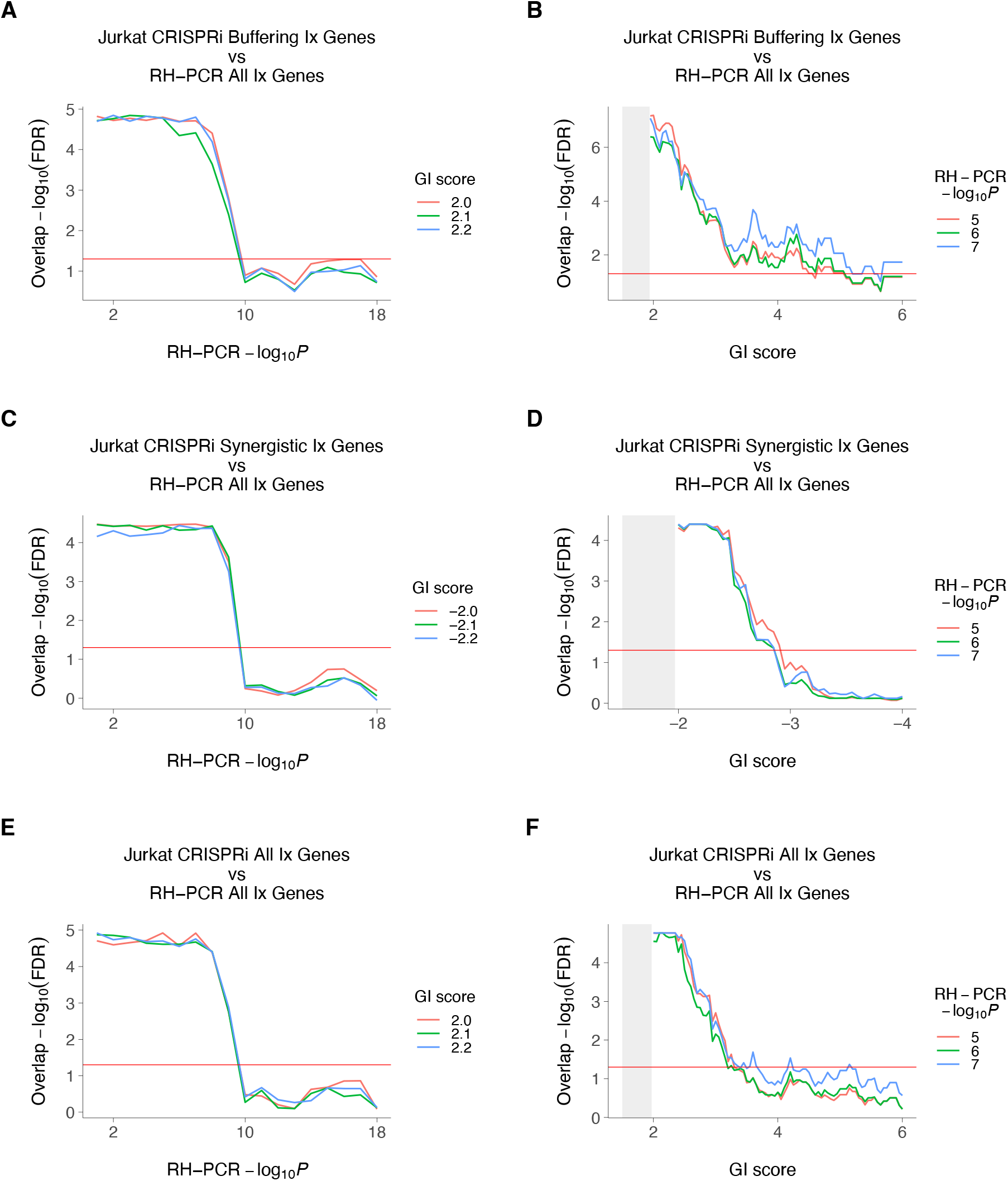
Genes in RH-PCR and Jurkat CRISPRi networks. *A,B*: Overlap of genes in Jurkat buffering CRISPRi and protein coding RH-PCR networks. *C,D*: Overlap of genes in Jurkat synergistic CRISPRi and protein coding RH-PCR networks. *E,F*: Overlap of genes in Jurkat combined buffering and synergistic CRISPRi and protein coding RH-PCR networks. FDR, false discovery rate. Gray rectangles at left of graphs in B, D and F indicate insignificant CRISPRi genetic interaction (GI) scores. Horizontal red lines, FDR (false discovery rate) = 0.05.

**Figure S10:**
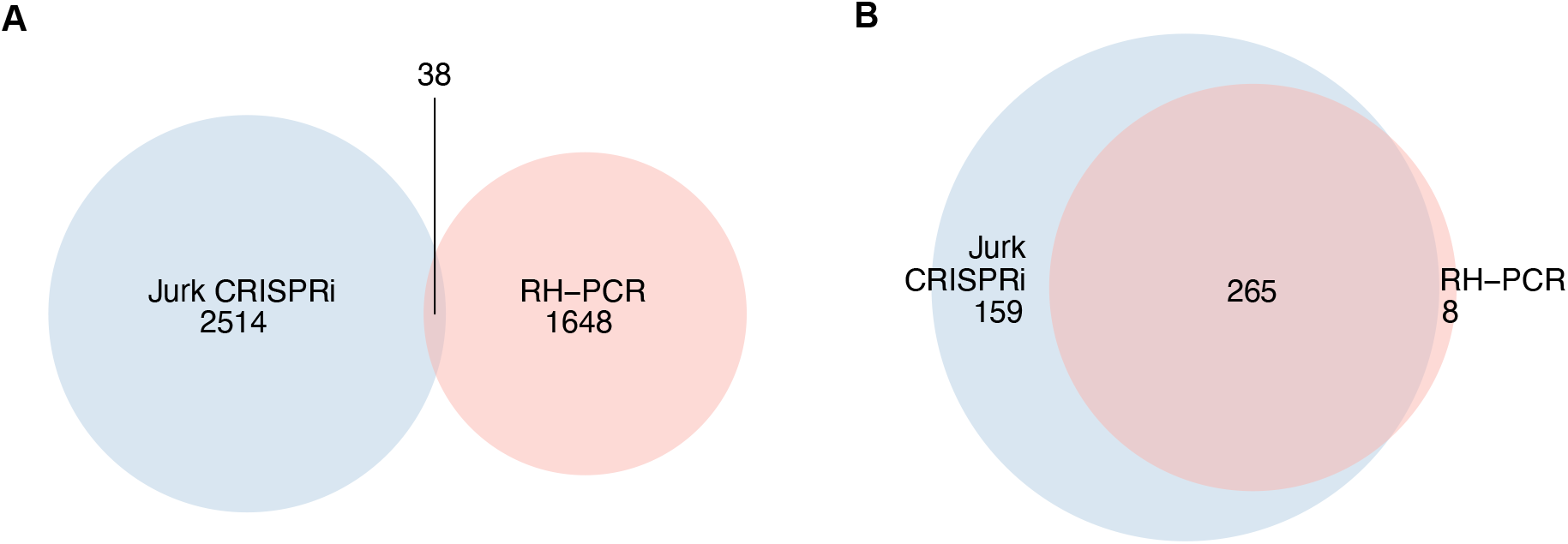
Interactions and genes in RH-PCR and Jurkat CRISPRi networks. *A*: Overlap of significant interactions in the essential RH-PCR and Jurkat combined buffering and synergistic CRISPRi networks. *B*: Overlap of genes participating in significant interactions in the two networks.

**Table S1:**
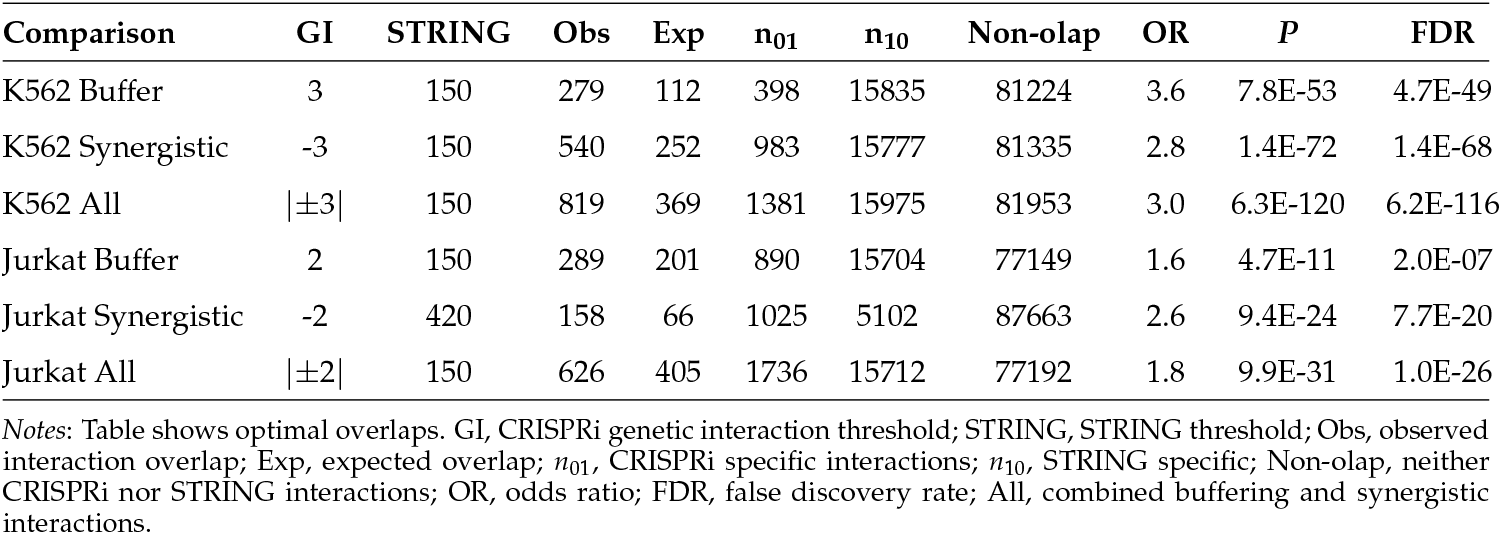
CRISPRi and STRING network overlaps.

**Table S2:** RH-PCR and STRING network overlaps.

| Comparison | -log<sub>10</sub>P | STRING | Obs | Exp | n<sub>01</sub> | n<sub>10</sub> | Non-olap | OR | P | FDR |
| --- | --- | --- | --- | --- | --- | --- | --- | --- | --- | --- |
| RH-PCR Ess | 2 | 150 | 400 | 285 | 1286 | 17229 | 85528 | 1.5 | 4.6E-13 | 4.7E-10 |
| RH-PCR ds | 1.5 | 160 | 3537 | 1738 | 62583 | 5239410 | 194184794 | 2.1 (0.03) | 1.5E-323 | 7.4E-321 |
| RH-PCR chrX | 3 | 170 | 3646 | 1822 | 83528 | 342478 | 16131323 | 2.1 | 2.1E-319 | 3.7E-316 |
| RH-PCR chrY | 1.5 | 160 | 91 | 36 | 5527 | 7874 | 1246547 | 2.6 | 3.8E-15 | 6.8E-12 |
| RH-PCR All | 4 | 770 | 4299 | 2255 | 1190102 | 372259 | 197923665 | 1.9 | 8.4E-323 | 4.3E-319 |

Notes: Table shows optimal overlaps. -log<sub>10</sub>P, RH-PCR significance threshold; STRING, STRING threshold score; Obs, observed interaction overlap; Exp, expected overlap; n<sub>01</sub>, RH-PCR specific interactions; n<sub>10</sub>, STRING specific; Non-olap, neither RH-PCR nor STRING interactions; OR, odds ratio; FDR, false discovery rate; RH-PCR Ess, RH-PCR network restricted to essential genes; RH-PCR ds, RH-PCR network randomly downsampled to same node number as essential genes; RH-PCR chrX, RH-PCR network downsampled so that each interaction has at least one gene on the X but not Y chromosome; RH-PCR chrY, RH-PCR network downsampled so that each interaction has at least one gene on the Y but not X chromosome; RH-PCR All, protein coding RH-PCR network. For the RH-PCR ds network, the standard deviation (s.d.) of the OR was evaluated using 20 independent downsamples at the optimal overlap.

**Table S3:**
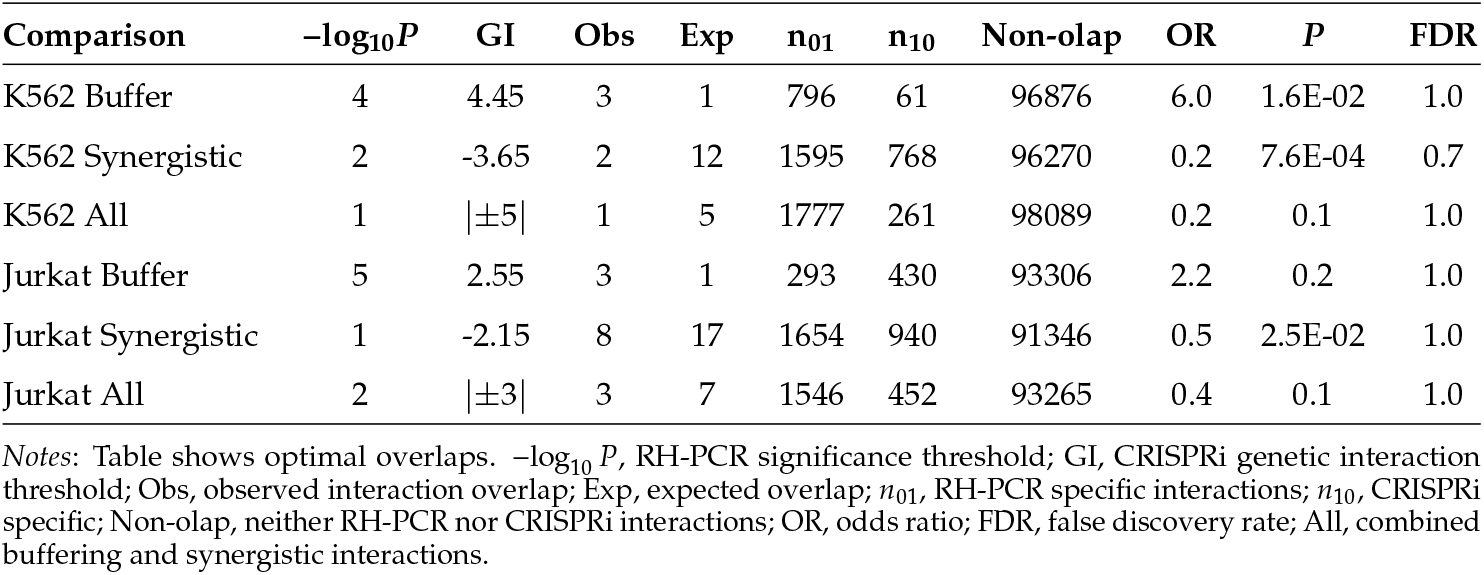
Essential RH-PCR network overlaps and CRISPRi.

**Table S4:**
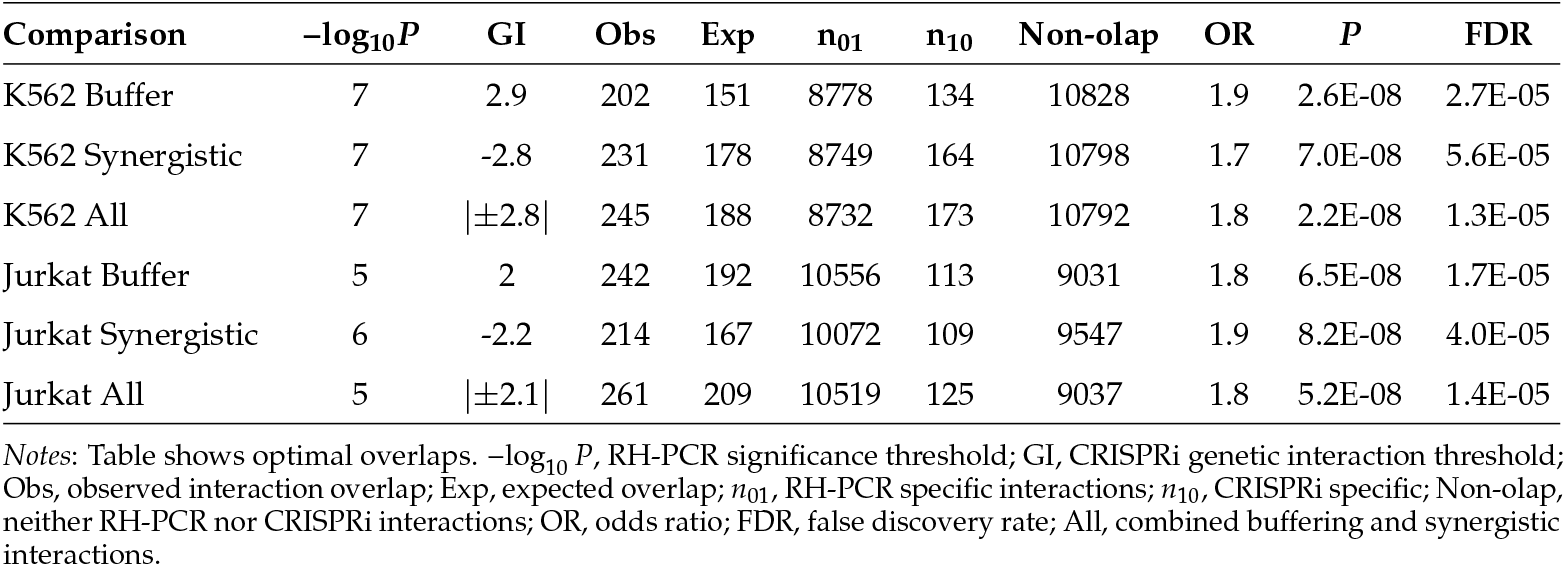
RH-PCR and CRISPRi overlaps of interacting genes.

| Comparison | $-\log_{10}P$ | GI | Obs | Exp | $n_{01}$ | $n_{10}$ | Non-olap | OR | $P$ | FDR |
| --- | --- | --- | --- | --- | --- | --- | --- | --- | --- | --- |
| K562 Buffer | 7 | 2.9 | 202 | 151 | 8778 | 134 | 10828 | 1.9 | 2.6E-08 | 2.7E-05 |
| K562 Synergistic | 7 | -2.8 | 231 | 178 | 8749 | 164 | 10798 | 1.7 | 7.0E-08 | 5.6E-05 |
| K562 All | 7 | $\vert \pm 2.8\vert $ | 245 | 188 | 8732 | 173 | 10792 | 1.8 | 2.2E-08 | 1.3E-05 |
| Jurkat Buffer | 5 | 2 | 242 | 192 | 10556 | 113 | 9031 | 1.8 | 6.5E-08 | 1.7E-05 |
| Jurkat Synergistic | 6 | -2.2 | 214 | 167 | 10072 | 109 | 9547 | 1.9 | 8.2E-08 | 4.0E-05 |
| Jurkat All | 5 | $\vert \pm 2.1\vert $ | 261 | 209 | 10519 | 125 | 9037 | 1.8 | 5.2E-08 | 1.4E-05 |

Notes: Table shows optimal overlaps. $-\log_{10}P$ , RH-PCR significance threshold; GI, CRISPRi genetic interaction threshold; Obs, observed interaction overlap; Exp, expected overlap; $n_{01}$ , RH-PCR specific interactions; $n_{10}$ , CRISPRi specific; Non-olap, neither RH-PCR nor CRISPRi interactions; OR, odds ratio; FDR, false discovery rate; All, combined buffering and synergistic interactions.

**Table S5:** RH-PCR and GWAS network overlaps.

| Comparison | $-\log_{10}P$ | GWAS | Obs | Exp | $n_{01}$ | $n_{10}$ | Non-olap | OR (s.d.) | $P$ | FDR |
| --- | --- | --- | --- | --- | --- | --- | --- | --- | --- | --- |
| RH-PCR chrX | 1.5 | 7 | 78 | 30 | 148846 | 3295 | 16408756 | 2.6 (0.7) | 3.2E-13 | 4.7E-10 |
| RH-PCR Essential | 1.5 | 5 | 210 | 114 | 1629 | 6260 | 96344 | 2.0 | 2.1E-17 | 1.9E-14 |
| RH-PCR Ess + 1 | 1.5 | 7 | 315 | 20 | 139866 | 11751 | 84185646 | 16.1 (2.6) | 9.7E-253 | 4.5E-250 |
| RH-PCR All | 4 | 9 | 6021 | 3525 | 1188380 | 582794 | 197713130 | 1.7 (0.1) | 1.1E-321 | 7.2E-319 |

Notes: Table shows optimal overlaps. $-\log_{10}P$ , RH-PCR significance value; GWAS, genome-wide association study $-\log_{10}P$ ; Obs, observed interaction overlap; Exp, expected overlap; $n_{01}$ , RH-PCR specific interactions; $n_{10}$ , GWAS specific; Non-olap, neither RH-PCR nor GWAS interactions; OR, odds ratio; FDR, false discovery rate; RH-PCR chrX, RH-PCR network downsampled so that each interaction has at least one gene on the X but not Y chromosome; RH-PCR Essential, RH-PCR network restricted to essential genes; Ess + 1, essential RH-PCR network plus genes linked by one interaction; RH-PCR All, protein coding RH-PCR network. For comparisons that employed a 20-fold downsample of the GWAS network, the standard deviation (s.d.) of the OR was evaluated using 20 independent downsamples at the optimal overlap.

**Table S6:** CRISPRi and GWAS network overlaps.

| Comparison | GI | GWAS | Obs | Exp | $n_{01}$ | $n_{10}$ | Non-olap | OR | $P$ | FDR |
| --- | --- | --- | --- | --- | --- | --- | --- | --- | --- | --- |
| K562 Buffer | 2.55 | 5 | 65 | 90 | 1410 | 5881 | 90380 | 0.7 | 6.0E-03 | 1.0 |
| K562 Synergistic | -8.7 | 11 | 5 | 0.7 | 16 | 3317 | 95297 | 9.0 | 5.6E-04 | 8.1E-02 |
| K562 All | $\vert \pm 8.7\vert $ | 11 | 5 | 0.7 | 16 | 3363 | 96744 | 9.0 | 5.6E-04 | 0.1 |
| Jurkat Buffer | 2.2 | 47 | 8 | 3 | 799 | 308 | 92917 | 3.0 | 6.4E-03 | 1.0 |
| Jurkat Synergistic | -5.8 | 5 | 6 | 2 | 21 | 6001 | 87920 | 4.2 | 6.3E-03 | 0.5 |
| Jurkat All | $\vert \pm 5.8\vert $ | 5 | 6 | 2 | 24 | 6085 | 89151 | 3.7 | 1.1E-02 | 0.5 |

Notes: Table shows optimal overlaps. GI, CRISPRi genetic interaction threshold; GWAS, genome-wide association study $-\log_{10} P$ threshold; Obs, observed interaction overlap; Exp, expected overlap; $n_{01}$ , CRISPRi specific interactions; $n_{10}$ , GWAS specific; Non-olap, neither CRISPRi nor GWAS interactions; OR, odds ratio; FDR, false discovery rate; All, combined buffering and synergistic interactions.

**Table S7:** RH-PCR protein coding network and GWAS subnetwork overlaps.

| Comparison | $-\log_{10} P$ | GWAS | Obs | Exp | $n_{01}$ | $n_{10}$ | Non-olap | OR (s.d.) | $P$ |
| --- | --- | --- | --- | --- | --- | --- | --- | --- | --- |
| Body mass index | 1.5 | 5 | 1983 | 1034 | 2710421 | 74057 | 196703864 | 1.9 | 9.7E-154 |
| Cancer | 1.5 | 5 | 8042 | 6959 | 2704362 | 503807 | 196274114 | 1.2 | 2.3E-37 |
| Colorectal cancer | 1.5 | 5 | 1491 | 1080 | 2710913 | 77941 | 196699980 | 1.4 | 1.0E-32 |
| Height | 1.5 | 5 | 90756 | 64465 | 2621648 | 4650464 | 192127457 | 1.4 (0.04) | 2.2E-308 |
| Smoking | 1.5 | 5 | 443 | 184 | 2711961 | 13093 | 196764828 | 2.5 | 9.9E-60 |

Notes: Table shows optimal overlaps. $-\log_{10} P$ , RH-PCR significance value; GWAS, genome-wide association study $-\log_{10} P$ ; Obs, observed interaction overlap; Exp, expected overlap; $n_{01}$ , RH-PCR specific interactions; $n_{10}$ , GWAS specific; Non-olap, neither RH-PCR nor GWAS interactions; OR, odds ratio. Full GWAS subnetworks employed except height, which used a two-fold downsampled subnetwork. The standard deviation (s.d.) of the height OR was evaluated using the two independent downsamples.

**Table S8:**
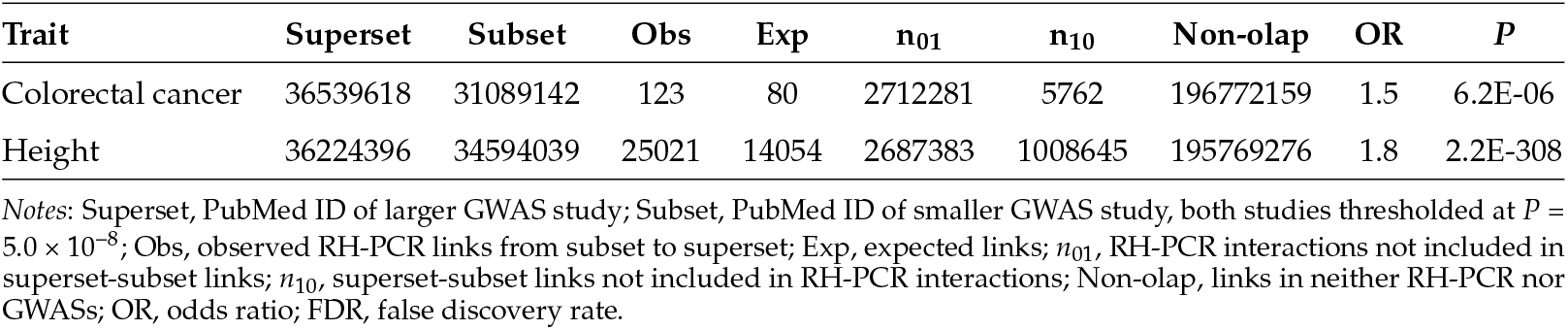
GWAS subnetwork overlaps based on RH-PCR network.

**Table S9:** CRISPRi, RH-PCR and STRING network overlaps using logistic regression.

| Comparison | GI | $-\log_{10} P$ | STRING | Z | $P$ | FDR |
| --- | --- | --- | --- | --- | --- | --- |
| K562 All | $\vert \pm 3\vert $ | – | 150 | 27 | 6.0E-156 | 5.5E-152 |
| Jurkat All | $\vert \pm 2\vert $ | – | 150 | 11 | 1.5E-30 | 1.4E-26 |
| RH-PCR Ess | – | 2 | 150 | 7.0 | 3.6E-12 | 2.7E-09 |
| RH-PCR All | – | 1 | 150 | 1649 | 2.2E-308 | 9.3E-308 |

Notes: Table shows optimal overlaps. GI, CRISPRi genetic interaction threshold; $-\log_{10} P$ , RH-PCR significance threshold; STRING, STRING threshold score; FDR, false discovery rate; K562 or Jurkat All, combined buffering and synergistic interactions; RH-PCR Ess, RH-PCR network restricted to essential genes; RH-PCR All, protein coding RH-PCR network.

